# MIC13-linked cristae disruption causes metabolic failure and early fibrotic remodelling in mitochondrial liver disease

**DOI:** 10.64898/2026.01.16.699229

**Authors:** Alexander Becker, Thomas O Eichmann, Kira an Mey, Viacheslav Vasiliev, Patrick Petzsch, Andrea Rossi, Felix Distelmaier, Ruchika Anand

## Abstract

Mitochondrial diseases are highly complex and heterogeneous, and nearly 20% of cases involve severe liver pathology. Here, we report an affected individual carrying a pathogenic *MIC13* variant (c.260-2A>G) associated with early-onset mitochondrial hepato-encephalopathy. Such mitochondrial hepatopathies are rare, multisystemic disorders with major liver involvement, difficult to diagnose, lack effective treatment, and are poorly understood, in part due to the absence of faithful disease-relevant cellular models. To investigate hepatocyte-specific consequences of the *MIC13* variant, we generated iPSCs carrying this disease-causing variant and differentiated them into induced hepatocytes (iHeps). MIC13, a key component of the MICOS complex required for cristae formation, was disrupted in these cells, and the resultant iHeps exhibited the same cristae defects observed in clinical samples. Integrated multi-omics and biochemical analyses revealed extensive metabolic rewiring, including disrupted amino acid turnover and accumulation of tricarboxylic acid (TCA) and urea cycle intermediates. Additionally, profound alterations in methionine cycle and transsulfuration pathways, along with enhanced bile acid synthesis, collectively affect methylation potential, redox homeostasis, and detoxification. Lipid metabolism was also impaired, with incomplete β-oxidation, increased ketogenesis, and diminished lipid storage. At the cellular level, extensive extracellular matrix (ECM) remodelling, increased intracellular collagen accumulation and enhanced cell migration indicated an early fibrotic phenotype. Overall, this clinically relevant model uncovers mechanistically how cristae defects drive metabolic imbalance and hepatocyte dysfunction, ultimately leading to early fibrotic changes in mitochondrial liver disease. These findings provide a strong mechanistic foundation for understanding mitochondrial liver disease and developing targeted therapeutic strategies.

## Introduction

Mitochondria play crucial roles in many cellular processes beyond energy conversion, including redox signalling, programmed cell death, and calcium homeostasis. Mitochondrial diseases are genetic disorders caused by pathogenic variants in mitochondrial or nuclear DNA, and often present as multi-organ or single-organ disease (1–3). Although neurological, cardiac, and skeletal muscle manifestations are most emphasized, an estimated 20% of mitochondrial diseases involve severe liver dysfunction (4–7). Liver is especially susceptible to mitochondrial dysfunction because of its central role in metabolic integration, detoxification and biosynthetic pathways. These disorders, termed mitochondrial hepatopathies often manifest as neonatal and childhood acute liver failure or, less commonly, as late-onset disease. Diagnosis is challenging because symptoms may mimic other liver or neuromuscular disorders, and liver transplantation is not possible due to multisystem involvement, especially the central nervous system (4).

Most implicated genes encode proteins involved in mitochondrial DNA (mtDNA) maintenance or electron transport chain (ETC) function, or even mitochondrial biogenesis and structure, including POLG, SCO1, MPV17, DGOUK, COX subunits and MIC13 (6). Mutations in the non-ETC gene *MIC13* (also known as *MICOS13* or *QIL1*) have been increasingly linked to early-onset mitochondrial hepato-encephalopathy (8–12). MIC13 is a key component of the mitochondrial contact site and cristae organizing system (MICOS) and essential for cristae and crista junctions (CJs) formation (13–17). Loss of MIC13 causes severe cristae abnormalities, including concentric-ring or stacked cristae membrane structures (14, 15). Although it is a very small protein (∼13 kDa) with no resemblance to any known protein so far, we previously identified conserved MIC13 residues that are crucial for its stability and function (18), demonstrated that MIC13 protects the MIC10-subcomplex from YME1L-mediated proteolysis (19), and identified SLP2 as a MIC13 interactor cooperatively regulating MICOS assembly and CJs formation (19). Despites these insights, the precise molecular functions of MIC13 and the mechanisms underlying the MIC13-associated pathology remain poorly understood.

MIC13-associated disease is a severe multisystem disorder that, in addition to neurological impairments including microcephaly and brain atrophy, and occasional pulmonary distress or nephrolithiasis, invariably presents with neonatal or infantile acute liver failure, often as the earliest clinical symptom (11). Histopathology commonly shows macro- or microvesicular steatosis, bile duct proliferation, enlarged hepatocytes with lipid-filled vacuoles, portal and perisinusoidal fibrosis, fibrotic septa, and reduced number of hepatocytes (12). Electron microscopy reveals marked disorganized cristae, often forming concentric ring-like structures (9, 10). Biochemical abnormalities include elevated lactate-to-pyruvate ratios, hypoglycaemia, coagulopathy, and increased urinary excretion of 3-methylglutaconic acid and tricarboxylic acid-cycle (TCA) intermediates. Affected individuals often do not survive beyond infancy or early childhood. Collectively, these reports implicate mitochondrial cristae integrity as an underappreciated determinant of hepatic metabolic homeostasis, however the mechanism by which MIC13-dependent cristae defects translate to hepatic metabolic failure remains unknown.

One of the major challenges for studying mitochondrial diseases is the lack of suitable model systems that faithfully mirror the disease mechanisms and tissue-specific phenotypes. Patient-derived fibroblasts and mouse models often fail to mimic these clinical features (20–22) and access to primary tissue, particularly paediatric liver, is limited. An alternative and increasingly powerful approach is to introduce pathogenic variants into genetically stable, well characterized induced pluripotent stem cells (iPSCs) (23–25). We recently reported the generation of the iPSCs harbouring MIC13 disease variant using CRISPR-Cas gene editing (26), enabling the investigation of MIC13-associated disease mechanism in a tissue-specific manner.

Here, we report an affected individual carrying a pathogenic *MIC13* variant, c.260-2A > G, who presented multisystem problems, including hepatopathy. In this study, iPSCs harbouring this disease-causing variant are differentiated into induced hepatocytes (iHeps), followed by integrated multi-omics and biochemical analyses to determine hepatocytes-specific pathological mechanisms. Our findings reveal that MIC13-mediated cristae defects and mitochondrial dysfunction drive extensive metabolic rewiring, including altered carbon, nitrogen, and lipid metabolism, together with extracellular matrix (ECM) remodelling and altered cell migration, collectively contributing to hepatocyte dysfunction and early fibrotic pathology in MIC13-associated disease.

## Results

### Clinical presentation of the *MIC13* c.260-2A > G variant

We report an affected individual carrying the *MIC13* c.260-2A > G variant presenting early neurological regression, treatment-refractory epilepsy, progressive hepatopathy and renal tubulopathy. This variant, also previously reported by Gödiker et al. as pathogenic in homozygous state, is located at the splice site upstream of exon 4 (10). It results in aberrant splicing, producing transcripts that cause either an in-frame deletion of seven amino acids or a frameshift with a premature stop codon, leading to loss of functional MIC13 protein. The previously reported individual presented with comparable phenotype including progressive encephalopathy, hepatopathy, and kidney disease (10).

### MIC13 deficiency does not influence the hepatic lineage specification in iPSCs

Given that MIC13-associated disease invariably presents liver dysfunction, we investigated the underlying mechanism by differentiating the recently generated iPSCs, harbouring the *MIC13* c.260-2A > G variant, to hepatocytes. These iPSCs retained pluripotency and exhibit complete loss of MIC13 due to the mutation (26).

The differentiation process involved the time-controlled addition of specific growth factors, mimicking the hepatocyte differentiation through intermediate stages including definitive endoderm, hepatic endoderm and induced hepatocytes (iHeps) (27, 28) (Fig 1A). Following the differentiation, iHeps visibly adopted a typical polygonal hepatocyte morphology distinct from the oval morphology of iPSCs (Fig 1A), indicating successful differentiation.

**Figure 1:**
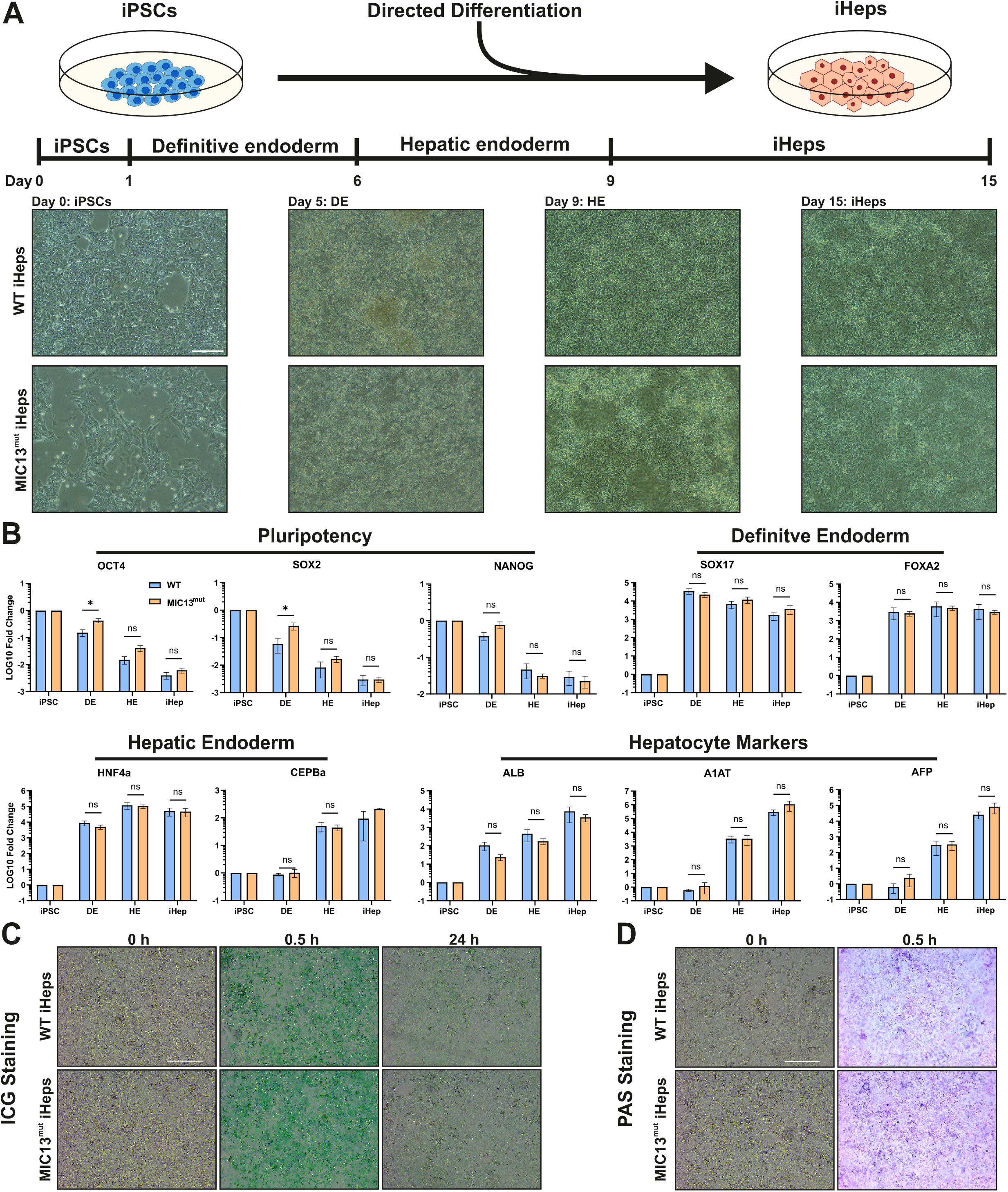
MIC13 is dispensable for hepatic lineage specification. (A) Schematic of the differentiation process with representative images of WT and MIC13^mut^ cells showing cellular morphology at different differentiation stages. Undifferentiated iPSC (at Day 0) display the typical oval or round shape, while differentiated induced hepatocytes (iHeps) (at Day 15) adopt a polygonal morphology, indicative of successful differentiation. Abbreviations: definitive endoderm (DE), hepatic endoderm (HE) and induced hepatocytes (iHeps). Scale bar: 100 μm (B) Quantitative PCR (qPCR) analysis was conducted at different differentiation stages using stage-specific gene markers: OCT4 (Octamer-binding transcription factor 4), SOX2 (Sex determining region Y-box 2), NANOG (Homeobox gene) for pluripotency in iPSCs; SOX17 (Sex determining region Y-box 17) and FOXA2 (Forkhead box 2) for definitive endoderm; HNFa (Hepatocyte Nuclear Factor a) and CEPBa (CCAAT Enhancer Binding Protein Alpha) for hepatic endoderm; and ALB (Albumin), A1AT (Alpha-1-antitrypsin) and AFP (Alpha-fetoprotein) for hepatocytes (iHeps). While there was a stage-specific decrease of pluripotency markers, iHeps markers increase steadily over the course of differentiation, peaking at iHeps stage. Expression of intermediate stage markers emerge at the corresponding stage of differentiation. This verifies the course of differentiation. Notably, all the markers (except OCT4, SOX2) are unaffected by the MIC13 deficiency across all stages of differentiation, demonstrating that MIC13 is not required for hepatic lineage acquisition. Data is showing log_10_ fold change of WT and MIC13^mut^ individually normalized to the ΔC_t_-values of the iPSC stage. It is represented as bar graph with mean ± SEM (n = 3 - 7). Statistical analysis was performed using Student’s *t* test. ∗*p*-value ≤0.05, ns = non-significant, *p*-value >0.05. HE marker CEPBa is exempt from statistical analysis at iHeps stage due to n = 2. (C) Functional assessment using indocyanine green (ICG) staining of iHeps shows uptake and release of the compound indicating intact metabolic activity in both WT and MIC13^mut^ iHeps. Scale bar: 100 μm. (D) Periodic acid-Schiff (PAS) staining demonstrates presence of glycogen storage in both WT and MIC13^mut^ iHeps. Scale bar: 100 μm.

To confirm the cell lineage, we performed quantitative PCR (qPCR) at various differentiation stages using stage-specific gene markers (Fig 1B). Pluripotency markers were highest in iPSCs and gradually decreased during differentiation, while the hepatic markers were strongly upregulated in iHeps, verifying lineage specification. Notably, the trajectory of differentiation was comparable between MIC13^mut^ and control iHeps (Fig 1B), indicating that MIC13 does not impair hepatic lineage acquisition.

To assess functional maturation of iHeps, we performed indocyanine green (ICG) uptake and release (Fig 1C) and Periodic acid-Schiff (PAS) staining (Fig 1D). Both cells efficiently took up ICG within 0.5 h and released it by 24 h, indicating efficient handling of the compound (Fig 1C). Strong PAS staining confirmed glycogen storage, supporting hepatocyte characteristics (Fig 1D). Together, these results show that despite the complete loss of MIC13, iHeps successfully acquire hepatic lineage and attain functional maturation.

### MIC13 deficiency causes mitochondrial and cristae morphology defects in iHeps

MIC13 is the key component of MICOS complex and its loss causes cristae defects and a reduction in the MIC10-subcomplex, which includes MIC10, MIC13, MIC26 and MIC27 (19). Western blot analysis of MIC13^mut^ iHeps confirmed a clear alteration in MIC10-subcomplex (Fig 2A), demonstrating the conserved role of MIC13 in stabilizing this subcomplex in hepatocytes (19). Transmission Electron Microscopy (TEM) analysis revealed abnormal cristae in MIC13^mut^ iHeps including stacked or concentric-ring-like cristae (Fig 2B), phenotype resembling the previously reported *MIC13* KO cells and the clinical samples (8, 14). While the total number of cristae remained unchanged, the number of CJs were drastically reduced in MIC13^mut^ iHeps (Fig 2C).

**Figure 2:**
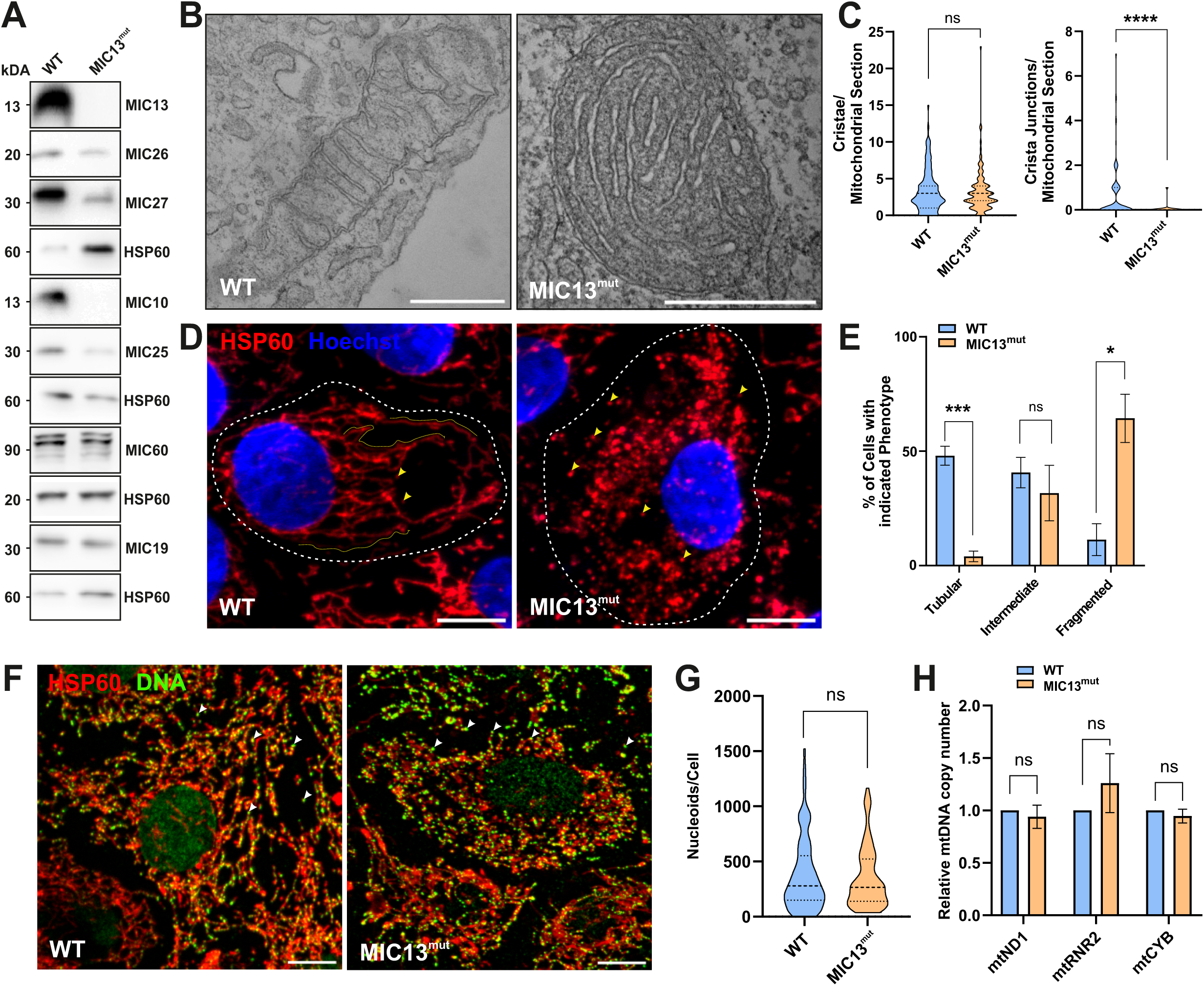
Loss of MIC13 disrupts the MIC10 subcomplex, crista junctions, and mitochondrial network morphology in iHeps. (A) Western blot analysis shows marked depletion of the MIC10-subcomplex (MIC10, MIC26, MIC27 and MIC13) in MIC13^mut^ iHeps, while the MIC60-subcomplex (MIC60, MIC19 and MIC25) remains largely unaffected, except for MIC25. (B) Representative TEM images of WT and MIC13^mut^ iHeps reveal abnormal cristae ultrastructure, including stacked or ring-like structure of the cristae. Scale bars: 500nm. (C) Quantitative analysis of TEM images show that while the number of cristae per mitochondrial sections remains unchanged between WT and MIC13^mut^ iHeps, there is a significant reduction in number of CJs in MIC13^mut^ iHeps, indicating impaired cristae organization. Data is shown as separate violin plot with range, median and interquartile values indicated from approximately 135 sections from three independent (n = 3) experiments. Statistical analysis was performed using Student’s *t* test. ∗∗∗∗*p*-value ≤0.0001, ns = non-significant, *p*-value >0.05. (D) Representative confocal images of WT and MIC13^mut^ iHeps immunostained for HSP60 (mitochondria, red) and counterstained with Hoechst (nuclei, blue) revealed extensive mitochondrial fragmentation in MIC13^mut^ iHeps. Yellow arrowhead indicate fragmented mitochondria and yellow lines traces tubular mitochondria. Scale bar: 10µm. (E) Quantification of mitochondrial network morphology, shown as a bar graph representing the percentage of cells displaying tubular, intermediate or fragmented mitochondria reveals a significant reduction in cells with tubular networks and a corresponding increase in cells with fragmented networks in MIC13^mut^ iHeps (around 50 cells per experiment, n = 3, mean ± SEM). Statistical analysis was performed using Student’s *t* test. ∗*p*-value ≤0.05, ∗∗∗*p*-value ≤0.001, ns = non-significant, *p*-value >0.05. (F) Representative confocal microscopy images of WT and MIC13^mut^ iHeps immunostained with anti-DNA (mitochondrial nucleoids and nuclei, green) and anti-HSP60 (mitochondria, red), reveal no detectable alterations in nucleoid distribution within mitochondria in MIC13^mut^ iHeps. White arrow heads indicate mtDNA nucleoids within mitochondria. Scale bar: 10µm. (G) Quantification of nucleoid count per cell shows no significant change between WT and MIC13^mut^ iHeps. Data is represented as a violin plot indicating the range, median and interquartile values, with data from around 90 cells from two independent experiments. Statistical analysis was performed using Student’s *t* test. ns = non-significant, *p*-value >0.05. (H) Quantitative PCR of genomic DNA shows no significant differences in the copy number of three mitochondrial genes, MT-ND1 (mitochondrially encoded NADH dehydrogenase 1), MT-RNR2 (mitochondrially encoded 16S RRNA), and MT-CYB (mitochondrially encoded cytochrome-b) in MIC13^mut^ iHeps. Data represented as bar graph (n = 3, mean ± SEM). Statistical analysis was performed using Student’s *t* test. ns = non-significant, *p*-value >0.05.

Analysis of overall mitochondria morphology showed highly fragmented mitochondria in MIC13^mut^ iHeps (Fig 2D, 2E), a phenotype associated with impaired bioenergetics and cellular stress (29). Although mtDNA depletions are observed in several mitochondrial hepatopathies (6), its role in MIC13-associated disease remains unclear, with only one report describing mtDNA loss in clinical samples (12). To investigate this, we assessed mitochondrial nucleoids using anti-DNA staining, which revealed no obvious change in the number of mtDNA foci (Fig 2F, 2G). qPCR with mtDNA-specific primers (mtND1, mtRNR2, and mtCYB) confirmed that mtDNA copy number remained unchanged across all loci tested (Fig 2H). These results indicate that observed mitochondrial defects in MIC13^mut^ iHeps arise independently of mtDNA depletion, primarily through cristae defects, although secondary or compensatory effects cannot be ruled out.

### MIC13 deficiency modifies the transcriptional and proteomic landscape in iHeps

To investigate the underlying mechanism driving MIC13-dependent changes in hepatocyte function, we performed a multi-omics analysis. Heatmap and Venn diagram visualizations of transcriptomics (RNA-seq) in iPSCs and iHeps derived from control and MIC13^mut^ display distinct cell-type specific transcriptional patterns, highlighting genes induced during hepatocyte differentiation and showing that MIC13^mut^ iHeps largely retain the capacity to acquire hepatic lineage identity (Fig 3A, 3B, supplementary Fig 1A). Focusing on iHeps-specific changes, we identified 1518 differentially expressed genes (DEGs) of which 600 were upregulated and 918 downregulated, and performed enrichment analysis both collectively for all the DEGs and separately for the up- or downregulated genes (Fig 3C, supplementary Fig 1B). Gene Ontology (GO) enrichment revealed that overall DEGs were associated with ECM organization, mitotic processes, cell migration, fatty acid and other monocarboxylic acid metabolic processes (Fig 3C). Notably, upregulated DEGs were predominately enriched in metabolic processes, particularly lipid and amino acid metabolism, while downregulated DEGs were mainly associated with cell cycle-related processes and ECM organization (Fig 3C). Consistent with this, canonical pathway analysis using ingenuity pathway analysis (IPA, Qiagen) identified Liver X Receptor/Retinoid X Receptor (LXR/RXR) activation, a key transcriptional regulator of lipid metabolism and inflammation (30), along with hepatic fibrosis and ECM organization among the top five enriched pathways, further highlighting dysregulation of metabolism and ECM in MIC13^mut^ iHeps (Table 1).

**Figure 3:**
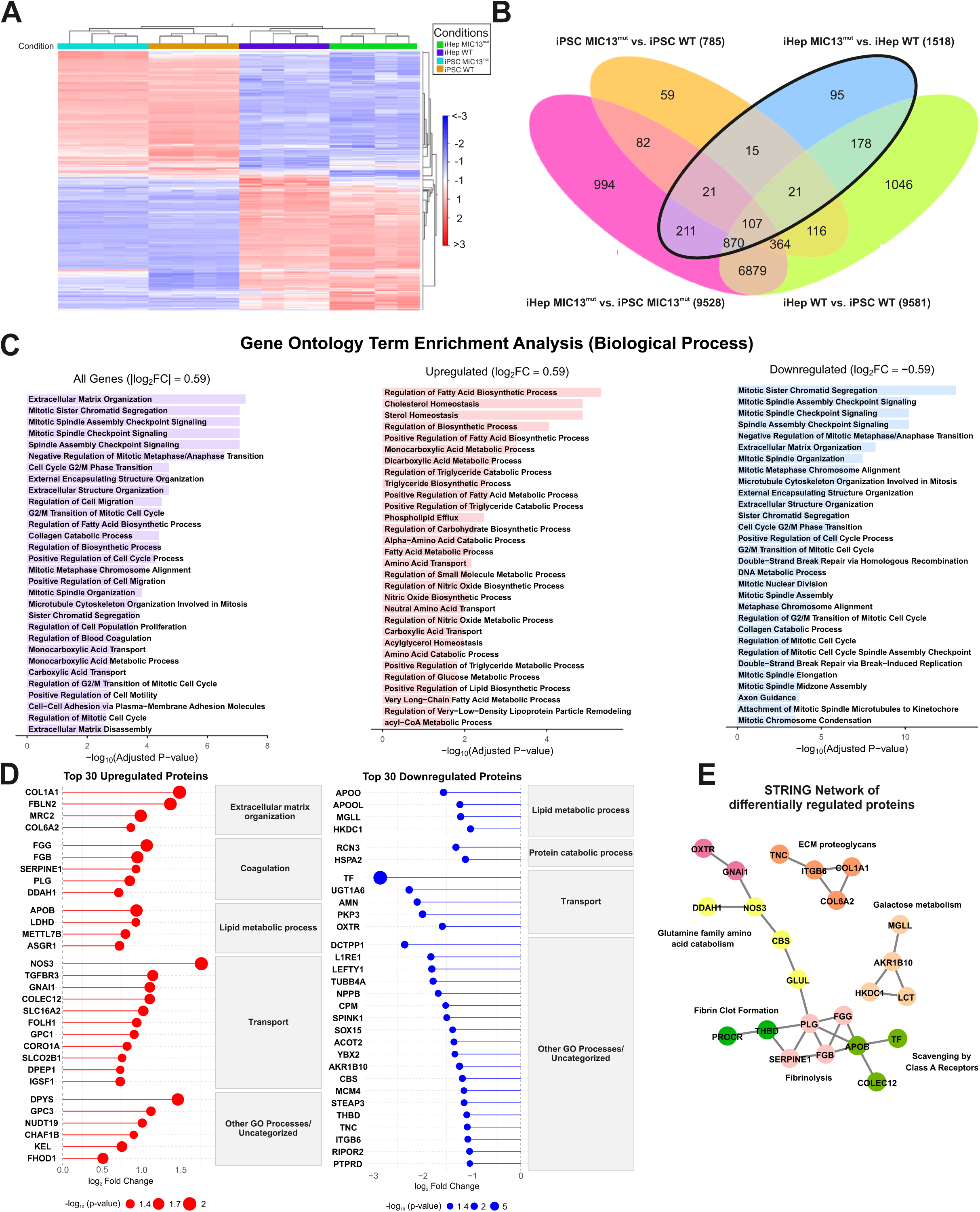
MIC13^mut^ iHeps display distinct changes in transcriptome and proteome. (A) Heatmap of differentially regulated genes (DEGs) in WT and MIC13^mut^ iPSCs and iHeps reveals strong transcriptional differences between cell types, confirming robust differentiation, while MIC13-dependent changes within each cell type were subtler and largely cell-type specific. (B) Venn diagram illustrating overlaps of DEGs across group comparisons. The iHep-specific differences between WT and MIC13^mut^ (bold outline) include 1518 DEGs, with 600 upregulated and 918 downregulated genes (Bonferroni-corrected *p* value, Fold change ≥ 1.5). (C) GO biological process term enrichment analysis in MIC13^mut^ iHeps was performed on all significant DEGs (|log_2_ Fold change| ≥ 0.59) as well as separately on upregulated (log_2_ Fold change > 0.59) and downregulated (log_2_ Fold change < −0.59). Analysis of all DEGs identifies pathways associated with extracellular matrix (ECM) organization, mitotic processes, cell migration and fatty acid metabolism. Upregulated genes are enriched for metabolic pathways, particularly lipid and amino acid metabolism, whereas downregulated genes are enriched for mitotic processes and ECM organization. (D) Proteomics analysis of MIC13^mut^ iHeps supports the transcriptomics trends. Among the top 30 upregulated proteins in MIC13^mut^ iHeps, GO-term based enrichment identifies ECM organization and coagulation as most enriched categories, whereas for the top 30 downregulated proteins, the highest categories correspond to lipid metabolic and protein catabolic processes. (E) STRING network analysis of significantly altered proteins in MIC13^mut^ iHeps reveals six distinct protein clusters, including Fibrinolysis, ECM proteoglycans, Fibrin clot formation, and Glutamine family amino acid catabolism, among others. Clusters are highlighted by distinct node colors.

**Table 1:** Top Canonical pathways identified by ingenuity pathway analysis (IPA, Qiagen) in MIC13^mut^ iHeps.

| Name | p-value |
| --- | --- |
| LXR/RXR Activation | 1,95E-17 |
| Hepatic Fibrosis / Hepatic Stellate Cell Activation | 1,03E-16 |
| Extracellular matrix organization | 1,48E-16 |
| Molecular Mechanisms of Cancer | 8,78E-16 |
| Sheddase Signaling Pathway | 2,25E-15 |

To confirm our observations, we next performed proteomics profiling on iHeps, which provided complementary insights into MIC13-dependent molecular changes. As the proteomics dataset contained a relatively small number of regulated candidates (∼97 proteins), large-scale enrichment analysis was not feasible. Therefore, we focused on the top 30 up- and downregulated proteins and grouped them according to the most representative GO biological process terms (Fig 3D). Many upregulated proteins were associated with ECM organization and coagulation, whereas the top downregulated proteins represented lipid metabolism and protein catabolism (Fig 3D), supporting key trends observed in our transcriptomics data. STRING interaction network of these proteins revealed distinct protein clusters such as ECM proteoglycans, fibril clot formation, glutamine family amino acid catabolism, among others (Fig 3E).

### Rewiring of central carbon and nitrogen metabolism in MIC13^mut^ iHeps

Hepatocyte mitochondria are central not only for energy conversion, but also for nitrogen disposal, redox balance and detoxification. Multi-omics analysis of MIC13^mut^ iHeps revealed substantial metabolic dysregulation, prompting us to perform untargeted metabolomics and integrate it with existing proteomics and transcriptomics data to directly assess the metabolic state. Classification of altered metabolites by chemical category revealed strong accumulation of dipeptides and amino acids in MIC13^mut^ iHeps (Fig 4A), indicating perturbed protein turnover and increased catabolic activity to meet energy and biosynthetic needs. Consistently, proteomics revealed altered expression of enzymes involved in protein turnover and autophagy, such as DPEP1, CPM and SQSTM1 (Supplementary Fig 2A).

**Figure 4:**
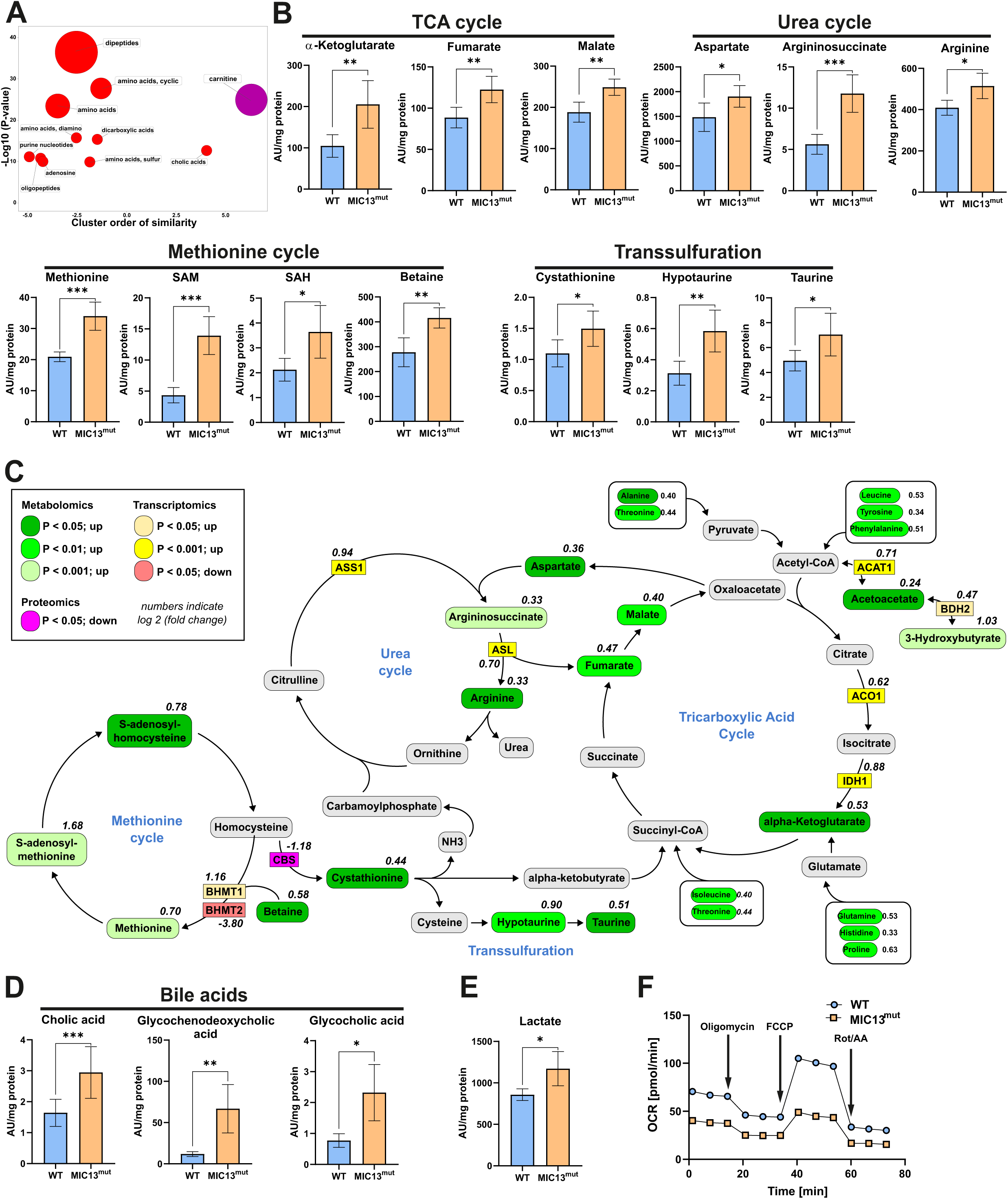
MIC13^mut^ iHeps exhibit broad rewiring of central carbon and nitrogen metabolism. (A) ChemRICH Plot of significantly altered metabolites in MIC13^mut^ iHeps reveals strong enrichment of dipeptides and amino acids, indicating broad change in amino acid metabolism and protein turnover. (B) In MIC13^mut^ iHeps, multiple metabolic pathways are altered. TCA cycle intermediates, including α-ketoglutarate, fumarate and malate, are significantly increased, suggesting altered TCA cycle activity. Urea cycle related metabolites, including aspartate, argininosuccinate and arginine, are significantly increased in MIC13^mut^ iHeps, reflecting altered nitrogen metabolism. Intermediates of methionine cycle, including methionine, S-adenosylmethionine (SAM), S-adenosylhomocysteine (SAH), and betaine were significant elevated in MIC13^mut^ iHeps, highlighting increased flux through methionine cycle and impaired methylation activity. Metabolites of the transsulfuration pathway, including cystathionine, hypotaurine, and taurine, are elevated, indicating enhanced flux towards glutathione biosynthesis and antioxidative capacity in response to redox stress in MIC13^mut^ iHeps. Data from five biological replicates. Statistical analysis was performed using Student’s *t* test. ∗*p*-value ≤0.05, ∗∗*p*-value ≤0.01, ∗∗∗*p*-value ≤0.001, ∗∗∗∗*p*-value ≤0.0001. (C) Schematic overview summarizing the metabolic rewiring of interconnected pathways in MIC13^mut^ iHeps. The diagram integrates metabolomic, transcriptomic and proteomic changes at key metabolic nodes, highlighting coordinated alterations across the TCA cycle, urea cycle, methionine cycle, and transsulfuration pathway. Elevated fumarate and aspartate link TCA and urea cycle flux, while homocysteine is diverted towards both remethylation and transsulfuration, reflecting simultaneous adjustments in methylation capacity and redox balance. Increased protein turnover provides amino acid that feed into the TCA cycle at multiple entry points. Additionally, the transcripts or proteins of key branch-point enzymes are altered, illustrating a coordinated transcriptional, proteomic and metabolic compensatory response to mitochondrial dysfunction. Abbreviations: ACO1 (Aconitase 1), IDH1 (Isocitrate dehydrogenase 1), ASL (Argininosuccinate lyase), ASS1 (Argininosuccinate synthase), BHMT1/2 (Betaine-homocysteine methyltransferase 1/2), CBS (Cystathionine B-synthase), ACAT1 (acetyl-CoA acetyltransferase 1), BDH2 (3-hydroxybutrate dehydrogenase 2). (D) Bile acid and its conjugates are increased, suggesting enhanced hepatic biosynthetic activity. (E) Lactate levels were elevated, indicating mitochondrial stress and increased glycolytic compensation. (F) Oxygen consumption rates (OCR) measured using Seahorse mitochondrial stress test reveals reduced mitochondrial respiration in MIC13^mut^ iHeps, consistent with widespread metabolic dysregulation and mitochondrial dysfunction. Representative traces from the complete experimental run is shown. (B, D, E) Data from five biological replicates. Statistical analysis was performed using Student’s *t* test. ∗*p*-value ≤0.05, ∗∗*p*-value ≤0.01, ∗∗∗*p*-value ≤0.001, ∗∗∗∗*p*-value ≤0.0001.

Several TCA intermediates like fumarate, malate and α-ketoglutarate were increased in MIC13^mut^ iHeps (Fig 4B), indicating altered central carbon metabolism and enhanced input from increased amino acids at multiple entry points. This is depicted in a pathway-level integration of metabolites, transcripts and proteins across affected metabolic pathways (Fig 4C). Fumarate, which links the TCA and urea cycle, and other urea cycle metabolites such as aspartate, arginine, and argininosuccinate were similarly elevated (Fig 4B), suggesting increased metabolic pressure on nitrogen disposal. Supporting this, transcripts of key branch-point enzymes (ACO1, IDH1, ASL, ASS1) were increased, indicating a coordinated transcriptional and metabolic rearrangement of the TCA and urea cycle regulation (Fig 4C).

In parallel, methionine, S-adenosylmethionine (SAM), S-adenosylhomocysteine (SAH), and betaine were elevated in MIC13^mut^ iHeps (Fig 4B), reflecting increased methionine cycle flux. Intermediates of transsulfuration, including cystathionine, hypotaurine and taurine were similarly increased (Fig 4B). In hepatocytes, the methionine cycle supports SAM-dependent methylation, whereas transsulfuration detoxifies homocysteine and contributes to glutathione production, maintaining redox balance. Homocysteine thus represents a key branch point, either remethylated to regenerate methionine or shunted into transsulfuration, as illustrated in our integrated pathway map (Fig 4C). The accumulation of intermediates of both pathways suggest a metabolic tension between sustaining the redox balance and meeting methylation demands. Accordingly, protein levels of CBS, a key enzyme of transsulfuration, were reduced, while transcript levels of BHMT1/2, which catalyse homocysteine remethylation, were altered (Fig 4C). This imbalance is further reflected in the accumulation of multiple SAM-dependent methylation products in MIC13^mut^ iHeps, (e.g increase in N-methylhistidine, trimethyllysine, methylnicotinamide, methylthioadenosine, along with decrease in guanidinoacetate) (Supplementary Fig 2B), indicating an overload of methylation capacity.

Regarding hepatocyte-specific functions, primary and conjugated bile acids were elevated (Fig 4D), consistent with the bile duct proliferation phenotype reported in affected individuals (9, 12). Lactate accumulation indicated mitochondrial stress and a shift toward glycolytic compensation (Fig 4E). Notably, fumarate, malate, lactate and methionine were elevated in several patient samples (9–11), and α-ketoglutarate was elevated in the patient described here, reinforcing the pathological significance of these metabolic changes as a systemic compensatory response to impaired mitochondrial function.

To determine whether these metabolic changes impaired mitochondrial respiration, we performed oxygen consumption rate (OCR) analysis using Seahorse Analyzer. MIC13^mut^ iHeps showed reduced mitochondrial basal respiration and other respiration-associated parameters, consistent with the broader metabolic stress (Fig 4F, supplementary Fig 2C). Altogether, these findings describe an interconnected metabolic phenotype in MIC13^mut^ iHeps, encompassing altered protein catabolism, TCA and urea cycle flux, methionine and transsulfuration pathways, redox and methylation imbalance, and disrupted bile acid metabolism, providing a mechanistic explanation for hepatocyte metabolic failure in MIC13-associated mitochondrial disease.

### Loss of MIC13 disrupts lipid metabolism, increases ketogenesis, and reduces lipid storage in iHeps

Another major outcome of the multi-omics analysis was a profound impairment in lipid metabolism. Hepatocytes are responsible not only for fatty acid uptake and oxidation, but also for triglyceride synthesis, transient lipid storage and systemic lipid homeostasis. Consistent with this, metabolomics revealed significant alterations in lipid handling in MIC13^mut^ iHeps, including increased long-chain fatty acids, reduced long-chain acylcarnitines, elevated short-chain acylcarnitines, accumulated ketone bodies (Fig 5A), along with changes in intermediates of phospholipids metabolism (supplementary Fig 3A). To understand the molecular basis of these changes, we examined lipid metabolism regulators in the proteomics and transcriptomics datasets. Proteomics revealed altered levels of key enzymes involved in fatty acid activation, trafficking and oxidation, including CPT1A, ACOT2, EHHADH, CROT, PLIN2 (Fig 5B), while transcriptomics revealed coordinated dysregulation of multiple fatty acid metabolism genes (Fig 5C, supplementary Fig 3B) in MIC13^mut^ iHeps, providing evidences for broad defects in fatty acid metabolism.

**Figure 5:**
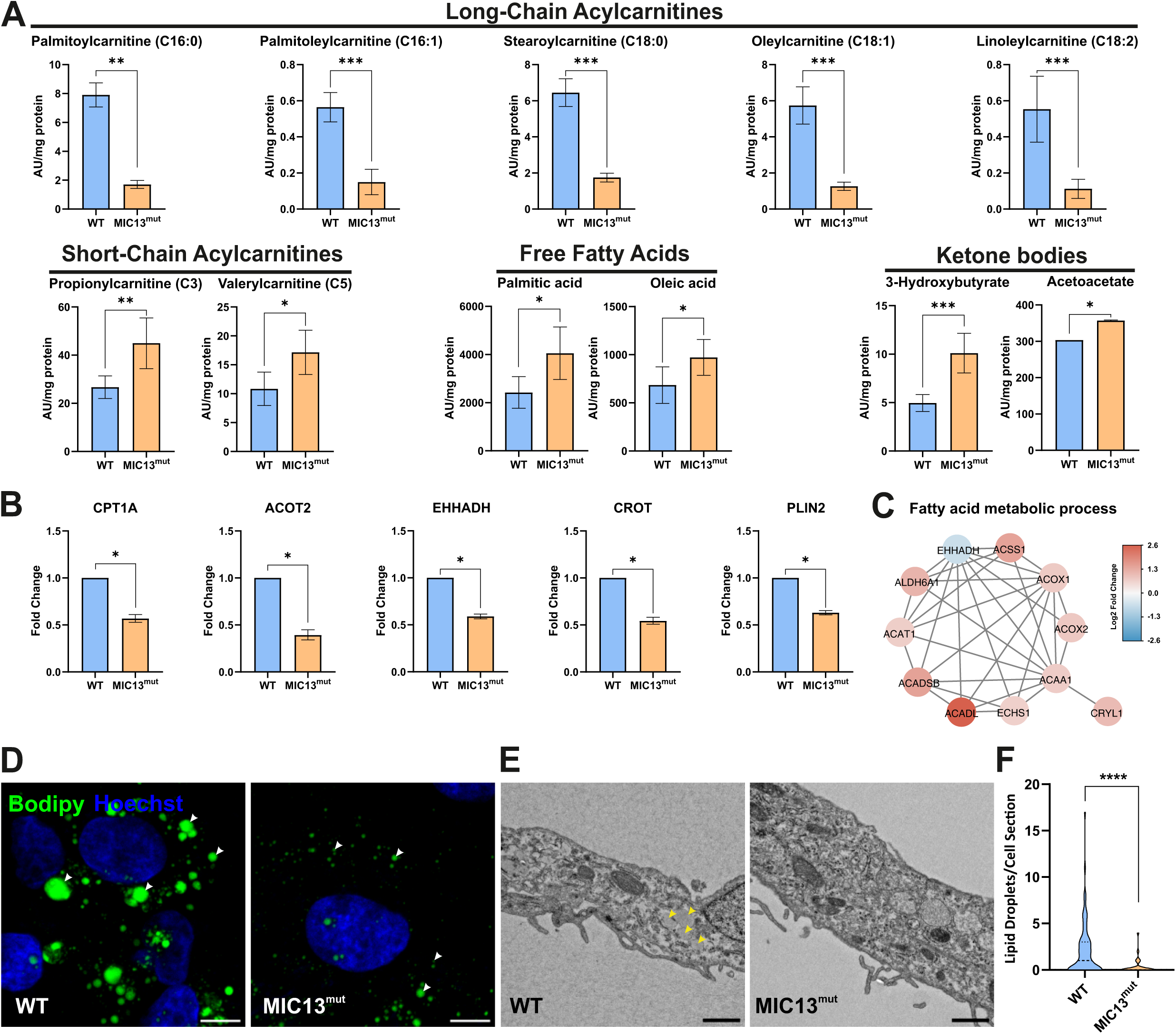
Loss of MIC13 impairs fatty acid metabolism, enhances ketogenesis and reduces lipid droplets. (A) In MIC13^mut^ iHeps, fatty acid metabolism is altered. Long-chain saturated and unsaturated acylcarnitines are significantly decreased, indicating impaired fatty acid activation and mitochondrial import. Conversely, short-chain acylcarnitines were increased, reflecting incomplete β-oxidation. Long-chain fatty acids, including palmitic and oleic acids, are elevated, consistent with impaired conversion to acylcarnitine. Ketone bodies are increased, suggesting a metabolic shift towards a compensatory response to altered TCA cycle flux and impaired fatty acid oxidation. Data from five biological replicates. Statistical analysis was performed using Student’s *t* test. ∗*p*-value ≤0.05, ∗∗*p*-value ≤0.01, ∗∗∗*p*-value ≤0.001, ∗∗∗∗*p*-value ≤0.0001. (B) Proteomics analysis shows significant downregulation of key proteins involved in fatty acid metabolism and lipid droplet biology, including CPT1A (Carnitine Palmitoyltransferase 1A), ACOT2 (Acyl-CoA Thioesterase 2), EHHADH (Enoyl-CoA Hydratase and 3-Hydroxyacyl CoA Dehydrogenase), CROT (Carnitine O-Octanoyltransferase) and PLIN2 (Perilipin 2), supporting broad defects in fatty acid activation, trafficking, oxidation and lipid storage. Data are represented as fold change relative to WT, error bars represent upper and lower limits calculated by back-transforming the log_2_ fold change ± SD. Data is collected from four biological replicates. (C) STRING network of DEGs from the MIC13^mut^ iHeps GO BP term enrichment as grouped by function. Specific genes show coordinated dysregulation, thus providing evidence of systemic lipid metabolic defects in MIC13^mut^ iHeps. Node colors represent the log_2_ fold changes. (D) BODIPY staining (green) of WT and MIC13^mut^ iHeps, counterstained with Hoechst (blue), shows marked reduction in lipid droplet accumulation in MIC13^mut^ iHeps. Arrow head marks lipid droplets. Scale bar: 10 μm. (E) Representative TEM images confirmed reduced lipid droplets in MIC13^mut^ iHeps. Lipid droplets were identified as round electron-lucent structures, marked by yellow arrow head. Scale bar: 1 μm. (F) Quantification of number of lipid droplets using TEM images demonstrates a significant reduction in lipid droplets abundance. Data are represented as violin plot where range, median and interquartile values are indicated. Data from around 80 sections from three independent experiments (n = 3). Statistical analysis was performed using Student’s *t* test. ∗∗∗∗*p*-value ≤0.0001

The increase in long-chain fatty acid such as oleic acid and palmitic acid, together with a decrease in their acylcarnitines and other long-chain acylcarnitines, suggests impaired conversion of fatty acids into acylcarnitines in MIC13^mut^ iHeps. Consistently, CPT1A, which mediates mitochondrial import of activated fatty acids for β-oxidation, was reduced. Lower levels of ACOT2, a key regulator of intramitochondrial acyl-CoA turnover, likely contributes to the accumulation of short-chain acylcarnitines, indicating incomplete β-oxidation. Reduced levels of peroxisomal enzymes EHHADH and CROT further indicate additional defects in peroxisomal β-oxidation, potentially limiting the supply of shortened fatty acid into mitochondria.

Despite impaired fatty acid oxidation, ketone bodies were markedly increased in MIC13^mut^ iHeps. This increase likely reflects altered TCA cycle together with increased amino acid-derived acetyl-CoA, shifting hepatocyte metabolism towards ketogenesis as a compensatory response (Fig 4C). The upregulation of the key ketogenesis enzymes, ACAT1 and BDH2, supports this shift (Fig 4C). Notably, elevated ketone bodies were consistently observed in patient samples, confirming the clinical relevance of this metabolic phenotype (12). Increased phospholipid metabolites further point to enhanced membrane remodelling under metabolic stress (supplementary Fig 3A).

These changes, especially reduction in PLIN2, a major lipid droplet coating protein, prompted us to analyse lipid droplets. We performed BODIPY staining and consistently observed reduced lipid droplets in MIC13^mut^ iHeps (Fig 5D). These results were confirmed by electron microscopy, where lipid droplets, identified as round electron lucent structures with poorly identifiable boundaries (31), were drastically reduced in MIC13^mut^ iHeps (Fig 5E, 5F). Overall, these finding reveal a coordinated collapse in fatty acid metabolism with impaired lipid storage and mobilization. Although livers of affected individuals show steatosis (12), detailed quantification and extent of lipid accumulation was not performed in the clinical samples. Our findings provide mechanistic insights into how the MIC13 pathological variant disrupts hepatocyte lipid homeostasis.

### MIC13 loss alters ECM homeostasis and enhances cell migration in iHeps

Having established that MIC13 deficiency in hepatocytes leads to profound mitochondrial, metabolic and lipid defects, we further examined whether these alterations affect cellular pathways governing hepatocyte function. In addition to metabolic pathways, transcriptomics analyses highlighted ECM regulation and cell cycle-related activities among the most significantly altered pathways. Because the livers of affected individuals display features such as fibrosis, bile duct proliferation, and general liver dysfunction, we investigated whether these cellular processes are reflected in our MIC13^mut^ iHeps model.

As a first step, we assessed whether MIC13^mut^ iHeps show abnormalities in cell cycle progression. FACS-based cell cycle profiling revealed no significant differences between WT and MIC13^mut^ iHeps across different cell cycle stages (supplementary Fig 4A, 4B). We therefore focused our subsequent analyses on ECM regulation, a hallmark of liver fibrosis and a key clinical feature of MIC13-associated disease (11, 32, 33). The ECM is a complex network of cross-linked macromolecules that provides structural support to tissues. While the liver has a high regenerative capacity, any chronic or persistent injury including metabolic or mitochondrial defects can disturb ECM homeostasis and wound healing responses and lead to pathological fibrosis (33, 34).

To define the nature of ECM alterations in MIC13^mut^ iHeps, we reassessed our multi-omics datasets in greater detail. Transcriptomics analyses revealed widespread dysregulation of genes involved in ECM homeostasis, affecting multiple major gene families and pathways (Fig 6A). These included structural ECM components (collagens, fibrillins, elastin, fibulins, fibronectins, nidogen-2), adhesion molecules and cell-surface receptors (integrin subunits), multiple matrix-remodelling enzymes (MMPs, ADAMs), key signalling components (TGFB2), and regulators of fibrillogenesis (LOXL1) (Fig 6A). STRING interaction network analysis of these ECM-related genes revealed distinct functional clusters involved in processes like collagen chain trimerization, elastin fibre formation, ECM disassembly, regulation of SMAD signalling and MMP activation (Fig 6B). Together, these analyses indicate disruption of multiple ECM organization, regulatory and signalling pathways in MIC13^mut^ iHeps. Proteomics further corroborated these findings, showing that many of the differentially abundant proteins were ECM-related including COL1A1, FBLN2, FGG, FGB, COL6A1, ITGB6, SPINK1, TNC, CCN1 (Fig 6C).

**Figure 6:**
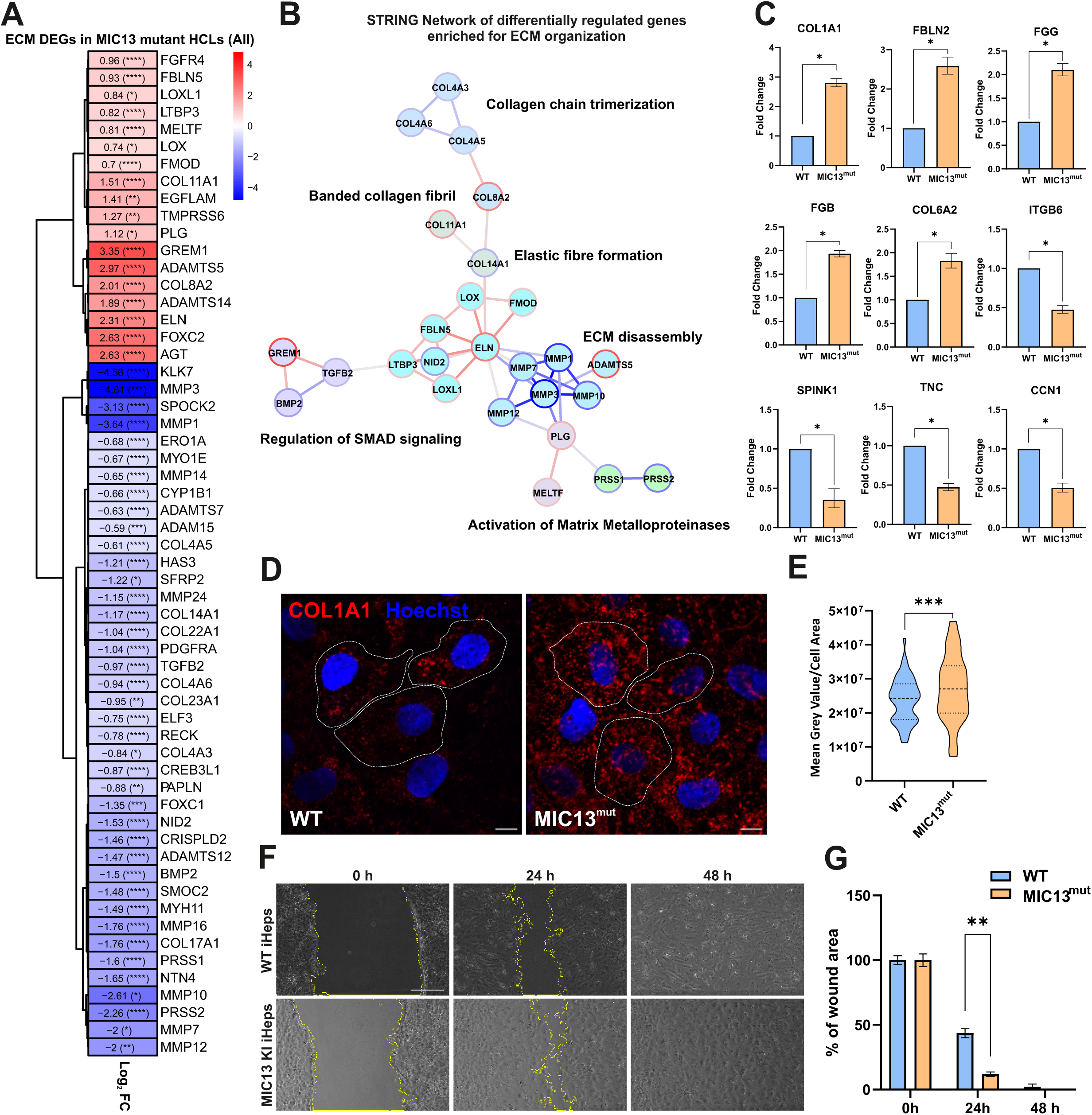
MIC13-dependent dysregulation of extracellular matrix and cell migration in iHeps. (A) Heatmap of differentially regulated genes (DEGs) in MIC13^mut^ iHep by GO term-based enrichment for ECM-related genes. DEGs were considered statistically significant with a cut-off fold change of ±1.5 and Bonferroni correction *p* ≤ 0.05. (B) STRING network of DEGs from the MIC13^mut^ iHeps GO BP term enrichment as grouped by function (colored nodes). The groups show various functions within extracellular matrix organization. Log_2_ fold changes are shown as a colored borders and edges of the respective nodes, revealing coordinative regulation (e.g. genes from the group “ECM disassembly” are mostly downregulated, whereas genes from “Elastic fiber formation” are upregulated. (C) Proteomics analysis of MIC13^mut^ iHeps shows that some of the most differentially altered proteins are ECM components, including COL1A1 (collagen type 1 alpha 1), FBLN2 (fibulin-2), FGG and FGB (fibrinogen gamma and beta chains), COL6A1 (collagen type V1 alpha 2), ITGB6 (integrin beta 6), Serine peptidase inhibitor, Kazal type 1 (SPINK1), TNC (tenasin C), and CCN1 (Cellular communication network factor 1). Data are represented as fold change relative to WT, error bars represent upper and lower limits calculated by back-transforming the log_2_ fold change ± SD. Data collected from four biological replicates. Statistical analysis was performed using Student’s *t* test. ∗*p*-value ≤0.05. (D) Immunofluorescence of COL1A1 (red) in iHeps with Hoechst nuclear counterstain (blue) reveals increased intracellular COL1A1 in MIC13^mut^ iHeps, suggesting either impaired secretion or biogenesis. Scale bar: 10μm. (E) Quantification of the COL1A1 signal shows a significant increase of COL1A1 abundance inside the MIC13^mut^ iHeps. Data is shown as violin plot with range, median and interquartile values indicated from approximately 100 cells analyzed from two independent experiments. Statistical analysis was performed using Student’s *t* test. ∗∗∗*p*-value ≤0.001. (F) Wound healing assay shows accelerated migration of MIC13^mut^ iHeps, as determined by wound closure over time, reflecting altered cell-ECM interactions. Scale bar: 100 μm (G) Quantification of the wound closure area demonstrates reduction in the area in MIC13^mut^ iHeps, indicating faster movement. Data are represented as mean ± SEM from three independent experiments. Statistical analysis was performed using Student’s *t* test. ∗∗*p*-value ≤0.01.

Given that COL1A1 is a key fibrotic marker and one of the most strongly altered protein in our proteomics analyses, we assessed its cellular localization using immunofluorescence.

COL1A1 exhibited a typical punctate staining in both WT and MIC13^mut^ iHeps (Fig 6D, 6E). However, MIC13^mut^ iHeps displayed remarkable increased fluorescence intensity compared to WT cells, indicating increased cellular abundance. This accumulation is consistent with altered collagen production or secretion and suggests early dysregulation of ECM homeostasis (32).

As ECM composition critically influences hepatocyte behaviour, including migration, we further assessed whether ECM alteration in MIC13^mut^ iHeps affects cell motility. Using wound healing assays, we observed that MIC13^mut^ iHeps migrated significantly faster than control cells, closing the wound area quicker (Fig 6F, 6G). This accelerated migration is consistent with altered cell-ECM interaction or disrupted signalling pathways. Together, these data support a model in which MIC13 deficiency, initiated by mitochondrial cristae defects, leads to systemic metabolic dysfunction that drives altered ECM remodelling and promotes aberrant hepatocyte migration, providing a pathological mechanism (Fig 7).

**Figure 7:**
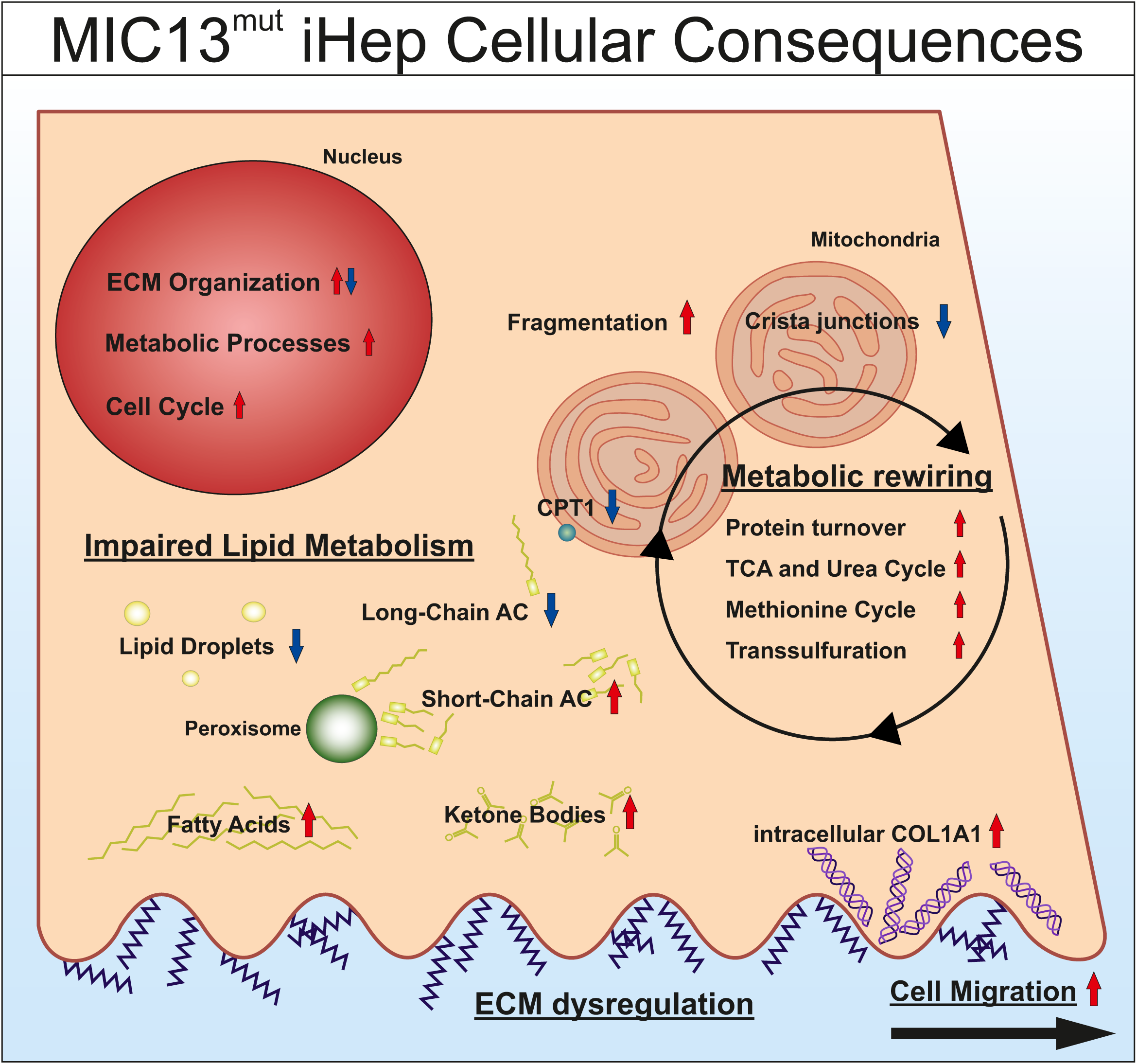
Proposed model for cellular consequences of MIC13 deficiency in iHeps. Loss of MIC13 leads to mitochondrial cristae defects without directly affecting the ETC or mtDNA maintenance, leading to extensive metabolic rewiring in hepatocytes. These changes include increased protein turnover and amino acid utilization, altered TCA cycle activity, disruption in urea cycle, methionine cycle, transsulfuration pathways, and impaired redox homeostasis and enhanced bile acid synthesis. Lipid metabolism was altered characterized by reduced lipid droplets and enhanced ketogenesis, decrease long-chain acylcarnitines (AC), and increased short-chain acylcarnitines and fatty acids, reflecting impaired fatty acid activation and oxidation. Reduced CPT1A and other fatty acid metabolism enzymes of mitochondria and peroxisomes indicate defective lipid handling. At the cellular levels, these metabolic disturbances were associated with altered expression of transcriptional programs related to ECM organization, mitosis, and metabolic regulation, leading to ECM remodeling, intracellular collagen accumulation and enhanced cell migration. Together, this model provides mechanistic insight into how systemic MIC13 deficiency drives mitochondrial liver disease, by linking primary cristae defects to metabolic failure and downstream cellular ECM alterations.

## Discussion

Overall, this study establishes an iPSC-derived hepatocyte model to investigate MIC13-associated mitochondrial hepatopathy. MIC13 is required for cristae organization and not directly involved in ETC or mtDNA maintenance. Nevertheless, its loss leads to extensive metabolic rewiring, including increased protein turnover, altered TCA cycle activity potentially fuelled by increased amino acids, and disruption in the urea cycle, methionine cycle, and transsulfuration, showing broad perturbations in carbon and nitrogen metabolism and redox homeostasis. These metabolic defects were accompanied by lipid metabolism alterations, including reduced lipid droplets and enhanced ketogenesis. These coincident metabolic and lipid changes likely drive broader alterations in cellular behaviour, including extensive ECM remodelling, increased intracellular accumulation of collagen and increased cell migration, features consistent with early events contributing to fibrosis (Fig 7). Together, our findings demonstrate that primary mitochondrial cristae defects are sufficient to drive profound hepatocyte metabolic and lipid rewiring, and ECM-related cellular changes (Fig 7), linking mitochondrial ultrastructure to early fibrotic processes in MIC13-associated hepatopathy.

Notably, liver-specific knockout of MICOS subunit *MIC19*, also causes hepatic alterations, highlighting general importance of cristae organization in liver function (35, 36).

Although iPSC-derived hepatocytes can retain fetal-like characteristics, our MIC13^mut^ iHeps recapitulates many clinical phenotypes, including accumulation of fumarate, malate and α-ketoglutarate, lactate, methionine, ketone bodies and other amino acids. The close concordance between clinical phenotypes and our findings support the physiological reliability of our pertinent cellular model and emphasizes the importance of tissue specificity and the careful choice of cell models in mitochondrial disease research (20–24).

Models of mitochondrial hepatopathies using iPSC, iHeps or organoid model are scarce, with only few available for DGUOK and POLG mutation (37–40). While these mtDNA depletion models revealed altered stress responses, such as ferroptotic sensitivity, which could be rescued by NAC and NAD (38, 40), it remains unclear whether different mitochondrial defects, including cristae disruption in MIC13 deficiency, elicit distinct or overlapping cellular responses. iPSC models also provide platform for therapeutic testing, including metabolic supplementation and identifying the mechanism of action for contraindicated drugs such as valproic acid (39).

Our metabolomics data revealed alterations across several interconnected metabolic pathways. While steady-state measurements provide a snapshot, future isotope tracing experiments using glucose, glutamine and lipid substrates could provide critical insights into the dynamics of these changes and define how MIC13 deficiency perturbs carbon, nitrogen and lipid metabolism (41, 42). The methionine cycle and transsulfuration pathways are particularly crucial for liver function, supporting methylation, redox balance and detoxification (43, 44). Its key mediator, SAM, supports epigenetic modifications, phospholipid synthesis, and detoxification reactions (45, 46). Dysregulation of SAM and related pathways has been implicated in the progression of liver diseases including steatosis, fibrosis and cirrhosis (44, 47), yet its significance in mitochondrial hepatopathies remains poorly understood. Further targeting this metabolic axis could provide therapeutic benefits, consistent with the reports that dietary methyl donors or choline supplementation improves liver function (48–50)

One of the most striking findings in our study was ECM dysregulation in MIC13^mut^ iHeps, yet how mitochondrial cristae defects contribute to ECM remodelling is poorly understood. ECM organization is highly energy-demanding (51), and mitochondrial dysfunction may limit ATP availability. Beyond energy supply, mitochondria influences ECM organization through multiple interconnected pathways (52), including provision of metabolites, ROS signalling and regulation of gene expression. For example, glutaminolysis drives collagen synthesis via α-ketoglutarate-dependent mTOR activation and proline hydroxylation (53), and in our dataset, increased glutamine, α-ketoglutarate and proline alongside collagen-related GO terms support this mechanistic link. Furthermore, excess ROS can modulate ECM remodelling by activating pro-fibrotic signalling pathways such as TGF-beta and HIF-1-α (54). Intracellular accumulation of COL1A1 could reflect altered production or processing, as mitochondrial stress perturbs ER-mitochondrial crosstalk, causing ER stress and defective procollagen processing (55). Metabolites such as SAM, α-ketoglutarate, and fumarate could also regulate chromatin-modifying enzymes governing ECM gene expression (56). Supporting this, liver-specific knockout of *MIC19* displayed enhanced collagen levels (36). Cell migration, another energy-intensive process, depends on reorganization of cytoskeletal and ECM components, and may be affected by mitochondrial dysfunction (57). While these links remain to be fully validated, few studies also highlight mitochondrial defects to ECM regulation, indicating this may be an underappreciated aspect in mitochondrial disease (52, 58, 59).

ECM regulation is complex and involves multiple liver cell types, each contributing differently to fibrosis (33). The defects we observed in hepatocytes may reflect mechanisms shared across cell types. Nevertheless, MIC13^mut^ iHeps provide a tractable and reproducible system to study ECM synthesis, organization, and turnover, and for guiding potential therapeutic strategies. While focused on a single MIC13 variant, the metabolic, lipid and ECM defects may extend to other mitochondrial hepatopathies. Moreover, our iPSC-derived platform provides a robust model to explore the broader multisystem consequences of these disorders.

## Material and Methods

### iPSC culture and maintenance

iPSCs were maintained in mTeSR™ Plus (STEMCELL Technologies) with 1% Penicillin/Streptomycin (P/S) (Sigma-Aldrich). Medium was replaced daily or every other day and cells were passaged upon reaching 75 % confluency on 12- or 6-well culture plates, coated with Geltrex™ LDEV-Free (1:100 dilution in DMEM/F-12, 37 °C for ≥ 1h). Prior to seeding, the 1x GelTrex solution was aspirated and iPSCs were seeded in mTeSR Plus medium containing 10 µM ROCK Inhibitor (Y-27632). Medium was changed the next day. For cell detachment, either ReLeSR™ (STEMCELL Technologies) was used for cell expansion, or Stempro™ Accutase™ Cell Dissociation Reagent (Gibco) for experiments that required cell counting. Before dissociations, cells were washed with DPBS (PAN-Biotech, without Ca^++^/Mg^++^) and incubated in ReLeSR or Accutase at 37 °C for 30 s and 5 min respectively. ReLeSR was aspirated to a thin film and the iPSCs were further incubated for 3-5 min before being resuspended in mTeSR and reseeded. On the other hand, Accutase was diluted with 10 mL DMEM/F-12 and the cells were collected and centrifuged at 200×g for 5 min. For cell counting (Neubauer haemocytometer), the Accutase-containing medium was removed and cells were resuspended in 1 mL mTeSR Plus with ROCK Inhibitor. For cryopreservation, 1×10^6^ cells, dissociated with Accutase, were pelleted at 300×g for 5 min, resuspended in 500 μL freezing medium (80 % FBS +20 % DMSO) containing 10 μM ROCK inhibitor, and stored at −80 °C before long-term storage in liquid nitrogen. Thawing involved rapid warming at 37 °C, dilution with complete medium, centrifugation (300×g, 5 min), and subsequent reseeding in medium containing 10 μM ROCK inhibitor.

### Differentiation of iPSCs to induced hepatocytes

One day before differentiation, 12-well culture plates were coated with Biolaminin 521 LN™ (BioLamina), diluted in ice-cold DPBS with Ca^++^/Mg^++^ to 5 μg/ml. Sealed plates were incubated overnight at 4 °C. Directed iHep differentiation was conducted as previously described (27, 60), with modifications to adapt the protocol to iPSC12 cell lines. Briefly, iPSCs were detached with Accutase as described above, and 3×10^5^ cells per well were seeded in mTeSR Plus with 10 µM ROCK Inhibitor in LN-521-coated 12-well culture plates (day 0). On the first day, differentiation medium was changed to definitive endoderm medium: 96% RPMI 1640 (PAN-Biotech), 2% B27 (without retinoic acid, Thermo Scientific), 1% Stable Glutamine (PAN-Biotech), 1% P/S (Sigma-Aldrich), 100 ng/mL Activin A (Peprotech), and for the first day, 2.5 μM CHIR99021 (Biomol). On day 6, definitive endoderm induction was completed and medium was replaced with hepatic endoderm medium: 76.5% KnockOut™ DMEM, 20% KnockOut™ Serum (both Thermo Scientific), 0.5% Stable Glutamine (PAN-Biotech), 1% P/S (Sigma-Aldrich), 0.01% β-Mercaptoethanol and 1% DMSO (both Carl Roth). On day 10, medium was changed to iHep medium for six days: 82% Leibovitz’ L-15 (PAN-Biotech), 8% FBS (Capricorn Scientific), 8% Tryptose Phosphate Broth (Thermo Scientific), 1% Stable Glutamine (PAN-Biotech), 1% P/S (Sigma-Aldrich), 1 μM human insulin (Santa Cruz), 10 ng/ml HGF (Peprotech), 20 ng/ml Oncostatin M (Immunotools) and 25 ng/ml dexamethasone (Sigma-Aldrich). Stage-specific medium was replaced daily throughout the differentiation protocol.

### Indocyanine green and Periodic acid-Schiff (PAS) staining

iHeps were stained with 1 mg/ml indocyanine green (Santa Cruz) directly in culture medium at standard culture conditions for 30 min and imaged before and after the staining using a light microscope (Leica, 40x). Then, the staining medium was aspirated and fresh iHep medium was added. Cells were cultured at standard culture conditions for 24 h and imaged again.

For PAS staining, iHeps were washed twice with pre-warmed DPBS without Ca^++^/Mg^++^ (PAN-Biotech) and fixed with 4% PFA at RT for 15 min. Cells were washed thrice with DPBS without Ca^++^/Mg^++^. Staining with the PAS kit (Sigma-Aldrich) was performed according to the manufacturer’s instructions. Briefly, 300 μl periodic acid solution was added per well and incubated at RT for 5 min. Subsequently, iHeps were washed thrice with 0.5 ml distilled water. Schiff’s reagent was added and incubated at RT for 15 min. Again, iHeps were washed thrice with 0.5 ml distilled water and counterstained with 300 μl solution per well and incubated for 90 s. iHeps were washed thrice with 0.5 ml distilled water before storage in DPBS and imaging with an inverted light microscope (Leica, 40x).

### Quantitative PCR

Total RNA was extracted from cells using the RNeasy Mini Kit (QIAGEN). Culture medium was aspirated and cells were washed with DPBS (PAN-Biotech). Subsequently, cells were lysed directly in culture well with RLT buffer and lysates homogenized by QIAshredder (QIAGEN) according to the manufacturer’s instructions. cDNA synthesis was carried out using the GoScript Reverse Transcription Mix (Promega) with 1 μg RNA. Similarly, gDNA was isolated with the Wizard Genomic DNA Purification Kit (Promega) according to the manufacturer’s instructions.

Quantitative real-time PCR was conducted using the Rotor-Gene 6000 (Corbett Research) with GoTaq qPCR Master Mix (Promega) according to manufacturer’s instructions. A list of all primers used is provided in supplementary Table 1. C_t_-values were normalized to the housekeeping gene RPS16. In the case of gene expression markers of differentiations, WT and MIC13^mut^ were individually normalized to the ΔC_t_-values of the iPSC stage. For genomic input, C_t_-values were normalized to the genomic housekeeping gene RPS16, MIC13^mut^ ΔC_t_-values were normalized to the ΔC_t_-values of the WT.

### Microscopy and quantification

For immunostaining, cells were washed twice with pre-warmed DPBS without Ca^++^/Mg^++^ (PAN-Biotech) and fixed with 4% PFA at RT for 15 min. The fixed cells were permeabilized by DPBS with 0.3% Triton-X (Sigma-Aldrich) and 3% BSA (Sigma-Aldrich) at RT for 15-45 min. Primary antibodies were diluted in DPBS and incubated with the cells at either RT for 1 h or overnight at 4 °C, in the dark. Subsequently, cells were washed thrice with DPBS and then incubated with secondary antibodies at RT for 30 min in the dark. Cells were washed thrice with DPBS and imaged using a spinning disk confocal microscope (PerkinElmer) with a 60x oil-immersion objective. Primary antibodies used were as follows: HSP60 (1:200, Sigma-Aldrich), COL1A1 (1:500, Abcam), Anti-DNA (1:100, Sigma-Aldrich). Secondary antibodies used were as follows: Alexa Fluor 568 Rabbit IgG, Alexa Fluor 488 Anti mouse (1:200, Life technologies).

To visualize lipid droplets, pre-warmed BODIPY staining solution, consisting of cell culture medium with 10 µM BODIPY 493/503 (Cayman Chemicals) was added directly to the cells and incubated at 37 °C for 30 min in the dark. Fixation was performed in the dark using 4% PFA. Imaging was conducted with a spinning disk confocal microscope (PerkinElmer) with an 60x oil-immersion objective.

Mitochondrial DNA nucleoids were quantified as previously described (61). Briefly, Z-stack images were captured using a spinning disk confocal microscope (PerkinElmer) with a 60x oil-immersion objective. In Fiji/ImageJ the image stacks were projected to one z-plane (Maximum projection) and 8-bit images manually thresholded (Otsu-Threshold). After using the watershed function, images were processed with a semi-automated CellProfiler4.2.5 pipeline. The number of mitochondrial nucleoids per cell was quantified.

### Electron microscopy

The control and MIC13^mut^ iHeps were fixed using 3% glutaraldehyde in 0.1 M sodium cacodylate buffer (pH 7.2). The cell pellets were rinsed in same cacodylate buffer and embedded in 2% agarose and stained using 1% osmium tetroxide for 50 minutes, followed by 1% uranyl acetate and 1% phosphotunstic acid solution for 1 h. Samples were dehydrated using a graded acetone series and embedded in Spurr epoxy resin, with polymerization at 65°C for 24 h. Ultrathin sections were prepared using an ultramicrotome, and images were acquired using TEM (Hitachi H7100 or JEM-2100 Plus, JEOL). Image analysis was performed in a double in a double-blind manner. Statistical analysis was performed using GraphPad Prism 8/10.

### SDS-PAGE and western blotting

The control and MIC13^mut^ iHeps were washed with ice-cold PBS, followed by protein extraction using RIPA lysis buffer. Protein concentrations were determined using Lowry assay (Bio-Rad, 5000113, 5000114, 5000115) and lysates were prepared in Laemmli sample buffer. Proteins were resolved on 10% or 15% SDS-PAGE and transferred into nitrocelluloase membrane (Amersham, 10600004), which was blocked for 1 h at room temperature (RT). The primary antibodies were incubated overnight at 4°C. The following antibodies were used: MIC10 (Abcam, 84969), MIC13 (custom made by Pineda (Berlin) against human MIC13 peptide CKAREYSKEGWEYVKARTK) (14), MIC19 (Proteintech, 25625-1-AP), MIC25 (Proteintech, 20639-1-AP), MIC26 (Thermo fisher Scientific, MA5-15493), MIC27 (Sigma-Aldrich, HPA000612-100UL), MIC60 (Abcam, ab110329), HSP60 (sigma, SAB4501464). Afterwards membranes were washed and incubated with HRP-conjugated secondary antibodies, either goat anti-mouse IgG (Abcam, ab97023) or goat anti-rabbit IgG (Dianova, 111-035-144). Chemiluminescent signals were detected using a Viber Lourmat Fusion SL imaging system (Peqlab).

### Analysis of cellular bioenergetics

The Seahorse XFe96 Analyzer (Agilent) and the Seahorse XF Cell Mito Stress Test assay (Agilent) were utilized to analyze iHep bioenergetics. Due to special handling requirements and continuous de-differentiation after detachment, the Mito Stress Test was standardized to meet these requirements, mainly by performing the assay shortly after cell detachment. XF sensor cartridges (Agilent) were hydrated overnight at 37 °C in a non-CO_2_ incubator. Cells were detached with Trypsin-EDTA (0,05 %, Capricorn Scientific) and 3.5×10^4^ iHeps per well were seeded in a Geltrex™-coated (BioLamina) Seahorse XF96 cell culture plate (Agilent) in Seahorse assay medium (5 mM D-galactose, 1 mM pyruvate and 2 mM L-glutamine, all PAN-Biotech). The plate was centrifuged at 200xg for 1 min and subsequently incubated for 1 h in a non-CO_2_ incubator. Mitochondrial oxygen consumption was measured after sequential addition of the following compounds: Oligomycin (1 μM), FCCP (1.25 μM) and rotenone/antimycin a (0.5 μM) according to the manufacturer’s instructions. Data was normalized to DNA amount as measured with Cyquant™ Cell Proliferation Assay Kit (Thermo Scientific).

### Wound healing assay

The wound healing assay was conducted with minor adaptations as described previously (62). Briefly, at the endpoint of iHep differentiation the 100% confluent cell monolayer was scratched in a straight line with a sterile 200 μl pipette tip. A designated spot along the scratch was imaged immediately (0h) and after 24 and 48 h using an inverted light microscope (Leica, 40x). Closure of the wound was quantified using Fiji/ImageJ, by measuring the area of the wound.

### Flow cytometry

DNA content was measured by propidium iodide (PI) staining. Cells were detached with Trypsin-EDTA (0,05 %, Capricorn Scientific) and centrifuged at 1000 x g for 5 min at 4 °C. iHeps were fixed by resuspending the cell pellet in 100 μl DPBS (PAN-Biotech) and drop-wise addition of 900 μl 80% ethanol. Cells were stored at -20 °C until sample preparation. Subsequently, cells were centrifuged at 500 x g for 10 min at 4 °C and supernatant discarded. Cells were incubated with 1mg/ml RNAse in DPBS for 30 min at RT. Supernatant was discarded and DNA stained by 50 μg/ml in DPBS on ice for 1 h. PI signal was measured using an Accuri™ C6 flow cytometer (BD Biosciences). At least 1×10^4^ cells were measured per condition analyzed. Data analysis was performed with FCS Express™ v7.24.0024.

### RNA-Seq Analyses

Sample preparations: Total RNA samples used for transcriptome analyses were quantified (Qubit RNA HS Assay, Thermo Fisher Scientific, MA, USA) and quality measured by capillary electrophoresis using the Fragment Analyzer and the ‘Total RNA Standard Sensitivity Assay’ (Agilent Technologies, Inc. Santa Clara, CA, USA). The samples in this study showed perfect RNA Quality Numbers (RQN) of 10.0. The library preparation was performed according to the manufacturer’s protocol using the ‘VAHTS™ Stranded mRNA-Seq Library Prep Kit’ for Illumina®. Briefly, 250 ng total RNA were used as input for mRNA capturing, fragmentation, the synthesis of cDNA, adapter ligation and library amplification. Bead purified libraries were normalized and finally sequenced on the NextSeq2000 system (Illumina Inc. San Diego, CA, USA) with a read setup of 1×100 bp. The Dragen BCL Tool (version 3.10.11) was used to convert the bcl files to fastq files as well for adapter trimming and demultiplexing.

Statistical Analysis: Data analyses on fastq files were conducted with CLC Genomics Workbench (version 23.0.2, Qiagen, Venlo, Netherlands). The reads of all probes were adapter and quality trimmed (using the default parameters). Mapping was done against the *Homo sapiens* (hg38; GRCh38.107) (July 20, 2022) genome sequence. After grouping of samples (four biological replicates) according to their respective experimental condition, the statistical differential expression was determined using the CLC Differential Expression for RNA-Seq tool (version 2.8, Qiagen, Venlo, Netherlands). The Resulting *p* values were corrected for multiple testing by FDR and Bonferroni-correction. A *p* value of ≤0.05 (*P*adj) was considered significant. The CLC Gene Set Enrichment Test (version 1.3, Qiagen, Venlo, Netherlands) was done with default parameters and based on the GO term ‘biological process’ (*H. sapiens*; June 22, 2022). The RNA-seq data was further evaluated using Ingenuity pathway analysis (IPA, Qiagen) with Content version: 153384343 (Release Date: 2025-11-21).

Data visualizations: All statistical visualization were performed in R environment (RStudio, Version 2025.09.2) (63) using the ggplot2 (64) for general graphics, ggrepel (65) for gene labelling, and pheatmap (66) for heatmap generation. A differential gene expression (DEG) analysis was conducted to identify and visualize significantly altered transcripts between the two genotypes. Pre-processed bioinformatics and statistical analyses data were imported to the R environment, and a Volcano Plot was generated using ggplot2 and ggrepel.

Gene Ontology (GO) biological processes (BP) enrichment analysis was performed using Enrichr (https://maayanlab.cloud/Enrichr/) (67–69) (last accessed on Dec. 16, 2025). Genes from the GO BP terms filtered for “extracellular matrix” were extracted and their log_2_ fold change were compiled. This filtered data was used as the input for a STRING interaction analysis and MCL Clustering (70). The STRING network was created using the interaction sources: Textmining, Experimental, Databases and Co-expression. The minimum interaction score was set to 0.700. MCL clustering was performed in STRING and the MCL cluster description as well as a short tabular output of the notes and edges were exported and used for reconstructing the network in Cytoscape (Version 3.10.4) (71).

### Proteomics

Data acquisition: Samples were analyzed by the CECAD Proteomics Facility on an Orbitrap Exploris 480 mass spectrometer equipped coupled to an Vanquish neo in direct flow setup (all Thermo Scientific). Samples were loaded onto a trap cartridge (#160434, Thermo Scientific) with a flow of 60 µl/min before reverse-flushed onto an in-house packed pulled tip column (30 cm length, 75 µm inner diameter., filled with 2.7 µm Poroshell EC120 C18, Agilent). Peptides were chromatographically separated with an initial flow rate of 400 nL/min and the following gradient: initial 4% B (0.1% formic acid in 80 % acetonitrile), up to 8 % in 3 min. Then, flow was reduced to 300 nl/min and B increased to 35% in 77 min, followed by column wash with 98% B at 400 nl/min and reequilibration to initial condition. MS1 scans were acquired from 390 m/z to 1010 m/z at 15k resolution. MS2 scans ranged from 400 m/z to 1000 m/z and were acquired at 15 k resolution with a maximum injection time of 22 ms, an AGC target of 1000%, and were acquired in 60 x 10 m/z windows with an overlap of 1 m/z. All scans were stored as centroid.

For the gas-phase fractionated library (72), a pool generated from all samples was analyzed in six individual runs covering the range from 400 m/z to 1000 m/z in 100 m/z increments using identical LC settings as the samples. For each run, MS1 was acquired at 60k resolution with a maximum injection time of 98 ms and an AGC target of 100%. MS2 spectra were acquired at 30k resolution with a maximum injection time of 60 ms. Spectra were acquired in staggered 4 m/z windows, resulting in nominal 2 m/z windows after deconvolution using ProteoWizard (73). Sample Processing in DIA-NN: The gas-phase fractionated library was build using DIA-NN 1.8.1 (74), a human Swissprot canonical database (UP5640, downloaded 15/01/2024) with settings matching acquisition parameters. DIA-NN was run with the additional command line prompts “—report-lib-info” and “—relaxed-prot-inf”. Further output settings were: filtered at 0.01 FDR, N-terminal methionine excision enabled, maximum number of missed cleavages set to 1, min peptide length set to 7, max peptide length set to 30, min precursor m/z set to 400, max precursor m/z set to 1000, cysteine carbamidomethylation enabled as a fixed modification. Afterwards, DIA-NN output was further filtered on library q-value and global q-value <= 0.01 and at least two unique peptides per protein using R (4.1.3). Finally, LFQ values calculated using the DIA-NN R-package. Afterwards, analysis of results was performed in Perseus 1.6.15 (75) by filtering for data completeness in replicate groups, FDR-controlled T-tests, and 1D enrichment analysis.

Visualization and analysis: Proteomic differential expression data was utilized to generate lollipop plots highlighting the top 30 most upregulated and top 30 most downregulated proteins in the MIC13^mut^ iHeps. These proteins were manually categorized into groups based on their Gene Ontology Biological Process (GO BP) terms. The plots were constructed using the ggplot2 package in R.

### Metabolomics

Cell pellets in 2 ml Safe-Lock PP-tubes were mixed with 400 µl cold MeOH (-20°C) and 100 µl cold ddH2O containing internal standards (IS, 4°C, alanine-13C2, serine-13C2, cholic acid-d4, octanoic acid-d15, glutarate, glycochenodeoxycholic aicd-d4, glycocholic acid-d4, Sigma-Aldrich or Biomol) and sonicated using a Bioruptor Pico (30 min, 4°C, 30sec ON/30sec OFF, frequency high; Diagenode). After addition of 300 µl ddH2O and 900 µl methyl tert-butyl ether (MTBE) samples were incubation under 1000 rpm shaking for 15 min on 4°C. After centrifugation (13,000 rpm, 10 min, 4°C) 800 µl of the upper phase were removed and replaced by 800 µl of artificial upper phase (from MTBE/MeOH/ddH2O, 9/4/4, v/v/v). After anew incubation and centrifugation, as described above, the complete upper phase was removed and 700 µl of the lower phase were collected, split (450 µl/250 µl) and dried using a vacuum concentrator. Metabolites (from 450 µl) were dissolved in 100 µl 70% acetonitrile (0.5 mM medronic acid) for further LC-MS analysis. Carboxylates (from 250 µl) were dissolved in 60 µl 75% MeOH for derivatization. Samples were derivatized by the addition of 30 µl 3 - Nitrophenylhydrazine (3NPH, 150 mM in MeOH), 30 µl 1-Ethyl-3-(3-dimethylaminopropyl)carbodiimide (EDC, 150 mM in MeOH), 30 µl pyridine (7.5% in 75% MeOH) and incubation for 30 min on ice followed by 30 min on 50°C under 800 rpm shaking. Samples were cooled down on ice, and the reaction was stopped by the addition of 50 µl formic acid (0.98%). Samples were centrifuged (13,000 rpm, 10 min, RT) and transferred to vials for further LC-MS analysis. For each analysis (metabolites and carboxylates), aliquots of all according samples were combined into a pooled quality control sample (QC) which was analyzed at the start and end of the sequence as well as after each fifth sample injection. An empty extraction (sample-free for background control) was injected before sample injections. An in-house reference compound mix (for level 1 identification) was injected at the start of the sequence. Each sample matrix-free injection was followed by two QC injections for column re-equilibration. Protein precipitates of the extraction were dried, solubilized in 250 µl NaOH (0.3 N, 55°C, > 4 h) and the protein content was determined using Pierce™ BCA reagent (Thermo Fisher Scientific) according to the manufacturer’s guidelines. Chromatographic separation was performed on a Vanquish UHPLC+ system (Thermo Fisher Scientific) equipped with an ACQUITY UPLC BEH Amide column (2.1 × 150 mm, 1.7 µm; Waters), using an 18 min gradient (metabolites negative mode, 400 µl/min) from 97% solvent A (ACN/ddH2O, 95/5, v/v; 10 mM NH4FA, 10 mM NH3) to 65% solvent B (ddH2O/ACN, 95/5, v/v; 20 mM NH4FA, 20 mM NH3) or a 23 min gradient (metabolites positive mode, 400µl/min) from 99% solvent A (ACN/ddH2O, 90/10, v/v; 10 mM NH4FA, 0.1% FA) to 50% B (ddH2O, 10 mM NH4FA, 0.1% FA) or a Hypersil Gold aq column (2.1 x 100 mm, 1.9 µm; Thermo) using a 30 min gradient (carboxylates negative mode, 400 µl/min) from 100% solvent A (ddH2O; 5 mM NH4FA, 0.1% FA) to 100% solvent B (ddH2O/ACN, 10/90, v/v; 5 mM NH4FA, 0.1% FA). The column compartment was kept at 40 °C, 30 °C and 40°C, respectively. A QExactive Focus mass spectrometer (Thermo Fisher Scientific) equipped with a heated electrospray ionization source (HESI II) was used for spectral acquisition. Metabolites were identified either level 1 via accurate m/z of the [M-H]- or [M+H]+ ion (<5ppm) and comparison of the retention time (rt) and MS2 spectra to synthetical reference compounds or level 2 without reference rt. All peaks (QC) were manually inspected in Freestyle (1.8 SP2) and peak extraction was performed in Skyline (24.1.0.199). Only metabolites with <25% peak area variation in QC samples were used for further processing. Peak areas were blank subtracted and normalized to IS and protein concentration. Metabolite data were expressed as AU (analyte-IS ratio)/mg protein.

For the Chemical Similarity Enrichment analysis of Metabolomics data (ChemRICH), a webtool has been used (https://chemrich.fiehnlab.ucdavis.edu/) (76).

### Statistical analysis

Data are represented as mean ± standard error of the mean (SEM). Statistical significance was determined by unpaired Student’s t-test. Data analysis was performed with Microsoft Excel. Data representation and statistical analysis was performed using GraphPad Prism8/10.

## Supporting information

all supplementary

## Acknowledgements

The research was supported by funding from Medical faculty of Heinrich Heine University Düsseldorf, FoKo-2020-71 (R.A., F.D. and A.R.), and Deutsche Forschungsgemeinschaft (DFG) grant, AN 1440/3-1 (R.A.), AN 1440/4-1 (R.A.), an Exploration grant from Boehringer Ingelheim Stiftung (R.A.) and a scholarship from Walter and Monika Neupert foundation to A.B. We thank Andrea Borchardt and Tanja Portugall for their excellent technical assistance with electron microscopy. Electron microscopy was performed at the Core facility for electron microscopy (CFEM) at the medical faculty of the Heinrich Heine University Düsseldorf. We also acknowledge the Centre for Information and Media Technology at Heinrich Heine University Düsseldorf for computational infrastructure and support. We are grateful to the CECAD (Cluster of Excellence for Aging Research) Proteomics facility in Cologne, Germany and especially to Dr. Stefan Müller and Dr. Prerana Wagle for support with the proteomics analyses. Proteomics facility work was supported by the large invest grant INST 216/1163-1 FUGG by the Deutsche Forschungsgemeinschaft. We also thank Prof. Gerhard Fritz and his student Ms. Sina Federmann for assistance with FACS experiments. We sincerely thank Prof. Andreas S. Reichert for guidance and discussions, and Dr. Arun Kumar Kondadi for continuous support and helpful input.

## Author Contribution statement

A.B.: Methodology, Validation, Formal Analysis, Investigation, Data Curation, Visualization, Writing-Original Draft, Writing-Review and Editing.

T.O.E.: Methodology, Formal Analysis, Investigation, Data Curation, Writing-Review and Editing.

K.A.M.: Methodology, Formal Analysis, Visualization.

V.V.: Formal Analysis, Data Curation Writing-Review and Editing. P.P.: Methodology, Formal Analysis, Investigation, Data Curation.

A.R. Funding Acquisition, Formal Analysis, Writing-Review and Editing.

F.D.: Conceptualization, Funding Acquisition, Formal Analysis, Data Curation, Writing-Review and Editing.

R.A.: Conceptualization, Funding Acquisition, Supervision, Project Administration, Formal Analysis, Data Curation, Writing-Original Draft, Writing-Review and Editing.

## Ethics statement

The study adheres to good clinical practice and follows the Declaration of Helsinki. Written informed consent for the pseudonymized use of clinical and genetic data of the unpublished MIC13 patient was obtained from the parents. Study for the iPSC-derived hepatocyte generation was approved by the local ethics committee of Heinrich Heine University Düsseldorf (Study number: 2021-1615).

## Data availability statement

The mass spectrometry proteomics data have been deposited to the ProteomeXchange Consortium via the PRIDE [1] partner repository with the dataset identifier PXD072091. The transcriptomics data is submitted to Gene Expression Omnibus (GEO) repository under the accession number GSE313628. All datasets will be made publicly available upon formal acceptance of the manuscript following peer review.

## Supplementary Information

### Supplementary Figure legends

**Supplementary Figure 1: Transcriptomics analysis of WT and MIC13^mut^ iPSCs and iHeps.**

(A) Principal component scatter plot showing the separation between WT and MIC13^mut^ iPSCs and iHeps.

(B) Volcano plots of differentially expressed genes (DEGs), the selected genes from the GO terms categories including ECM organization, fatty acid metabolism, lipid metabolism are indicated.

**Supplementary Figure 2: Metabolic alterations in MIC13^mut^ iHeps**

(A) Proteomics analysis reveals significantly altered proteins belonging to protein metabolism and autophagy, DPEP1 (dipeptidase 1), CPM (carboxypeptidase M) and SQSTM1 (sequestosome-1). Data collected from four biological replicates. Statistical analysis was performed by Student’s *t*-test with *: *p* value < 0.05.

(B) Metabolomics data reveals significant changes in the SAM-dependent methylation metabolites, indicating impaired methylation in MIC13^mut^ iHeps. Data collected from five biological replicates. Statistical analysis was performed by Student’s *t*-test with *: *p* value < 0.05, ***: *p* value < 0.001.

(C) Bar graph representing the oxygen consumption rates (OCR), including BR (basal respiration), MR (maximal respiration), SRC (spare respiratory capacity), PL (protein leak), ATP prod (ATP production) and NMOC (non-mitochondrial respiration) from four independent Seahorse Analyzer runs, indicate a trend towards reduce respiration in MIC13^mut^ iHeps.

**Supplementary Figure 3: Alterations associated with lipid metabolism in MIC13^mut^ iHeps**

(A) Metabolites related to phospholipid metabolism were significantly altered in MIC13^mut^ iHeps. Data collected from five biological replicates. Statistical analysis was performed by Student’s *t*-test with *: *p* value < 0.05, **: *p* value < 0.01, ***: *p* value < 0.001.

(B) Transcriptomics data analysis reveals significant changes in genes associated with fatty acid metabolism in MIC13^mut^ iHeps. Statistical analysis was performed by Student’s *t*-test with ***: *p* value < 0.001.

**Supplementary Figure 4: Cell-cycle progression in MIC13^mut^ iHeps using FACS-based assay.**

(B) FACS-based analysis showed no significant difference in distribution of the of cells across different cell-cycle phases between WT and MIC13^mut^ iHeps (n = 3).

(C) Representative FACS gating profile used to determine cell-cycle phases in WT and MIC13^mut^ iHeps.

**Supplementary table 1:**
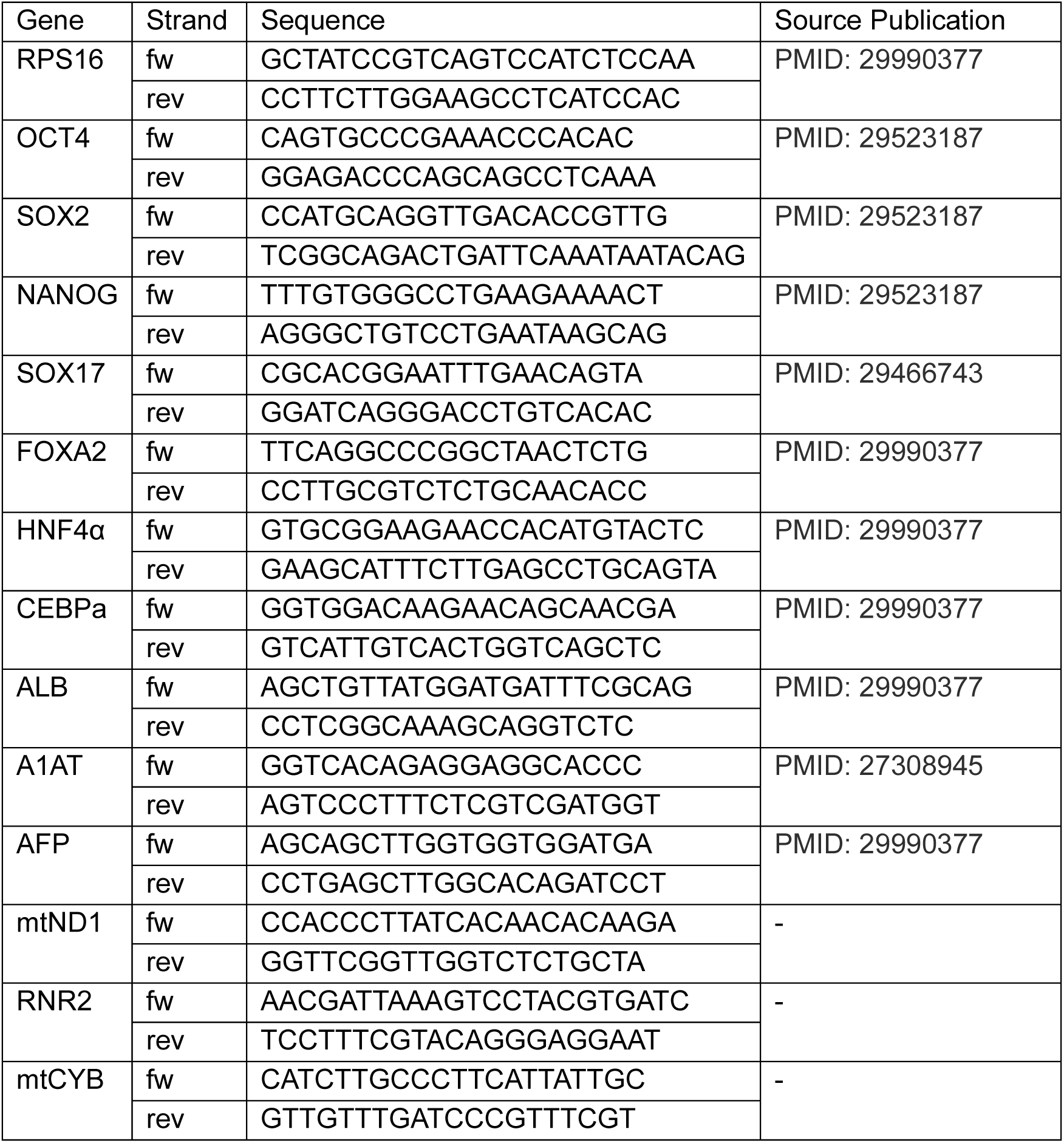
Sequences of primers used in this study.

