## Supplementary material for "MIC13-linked cristae disruption causes metabolic failure and early fibrotic remodelling in mitochondrial liver disease": all supplementary

### Supplementary Figure 1

A

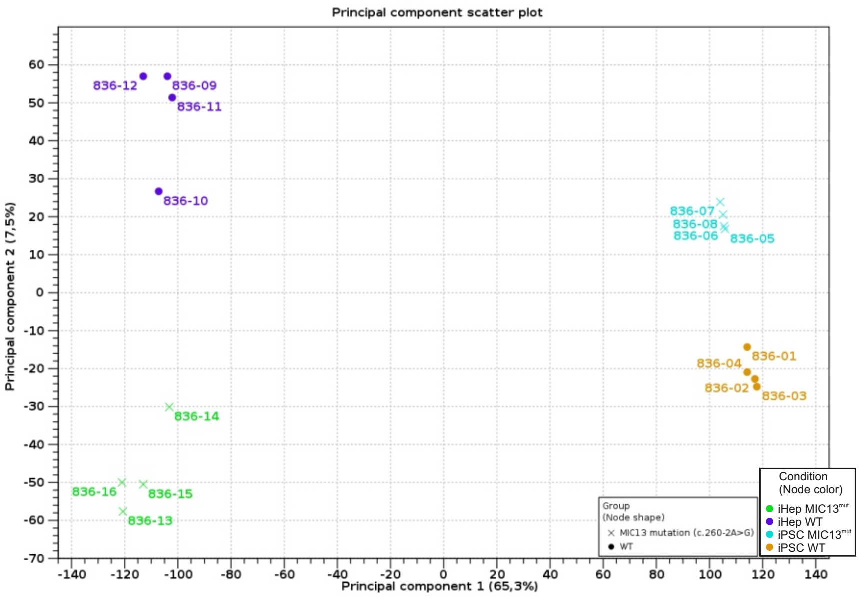


B

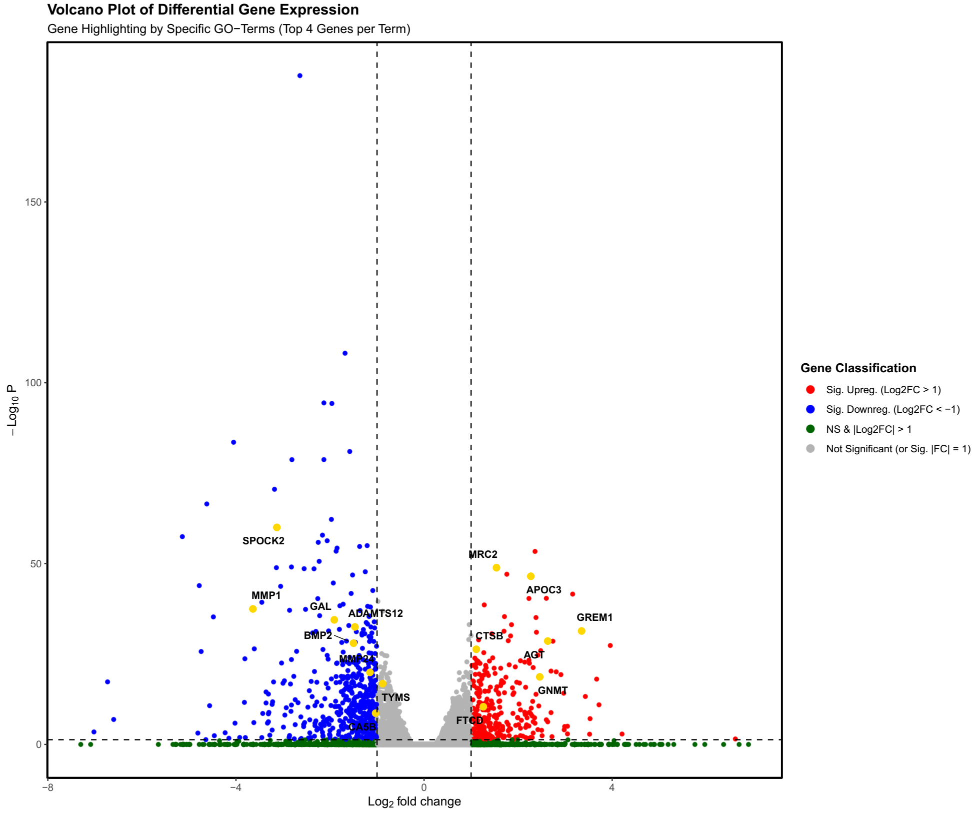

### Supplementary Figure 2

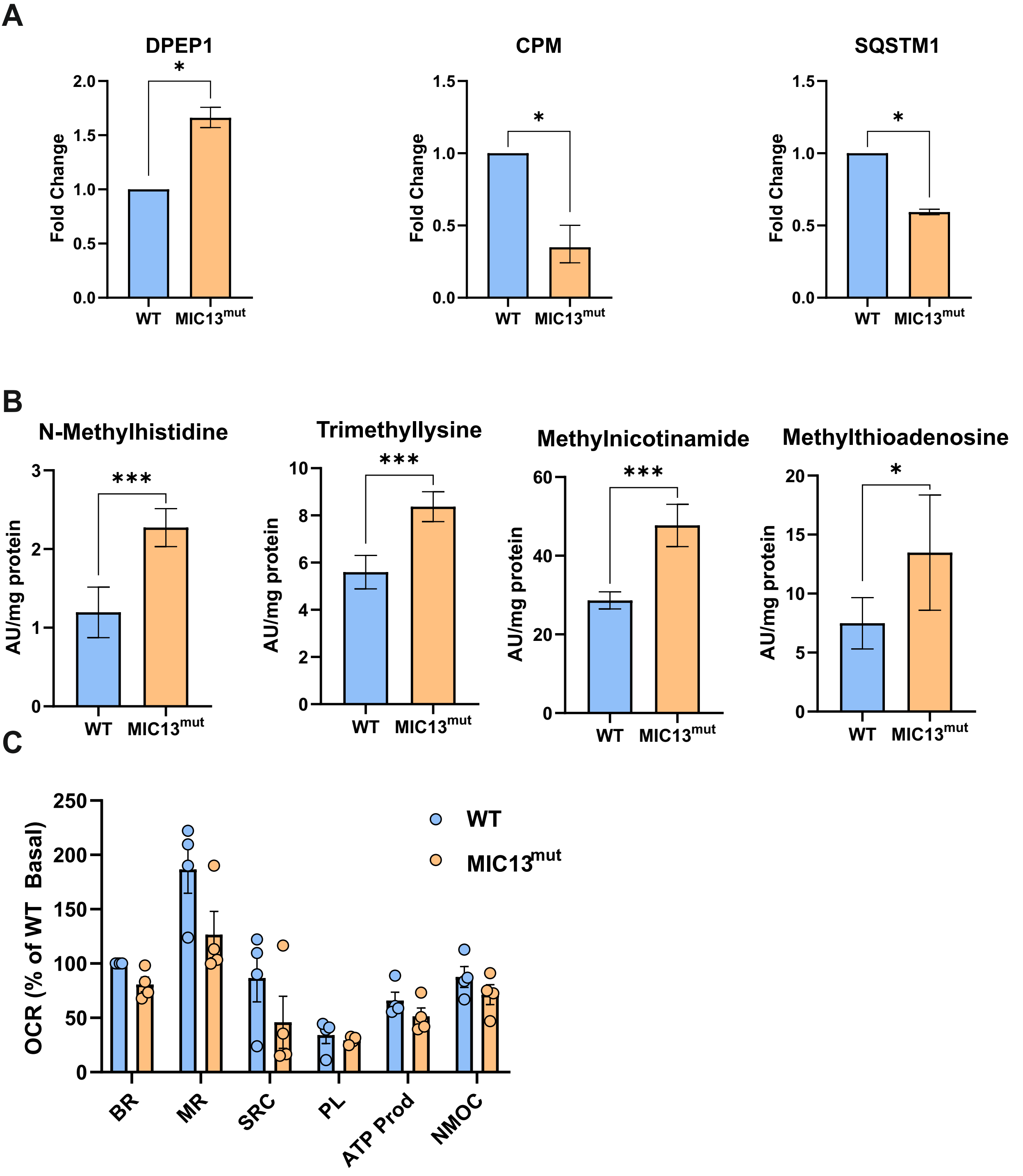

### Supplementary Figure 3

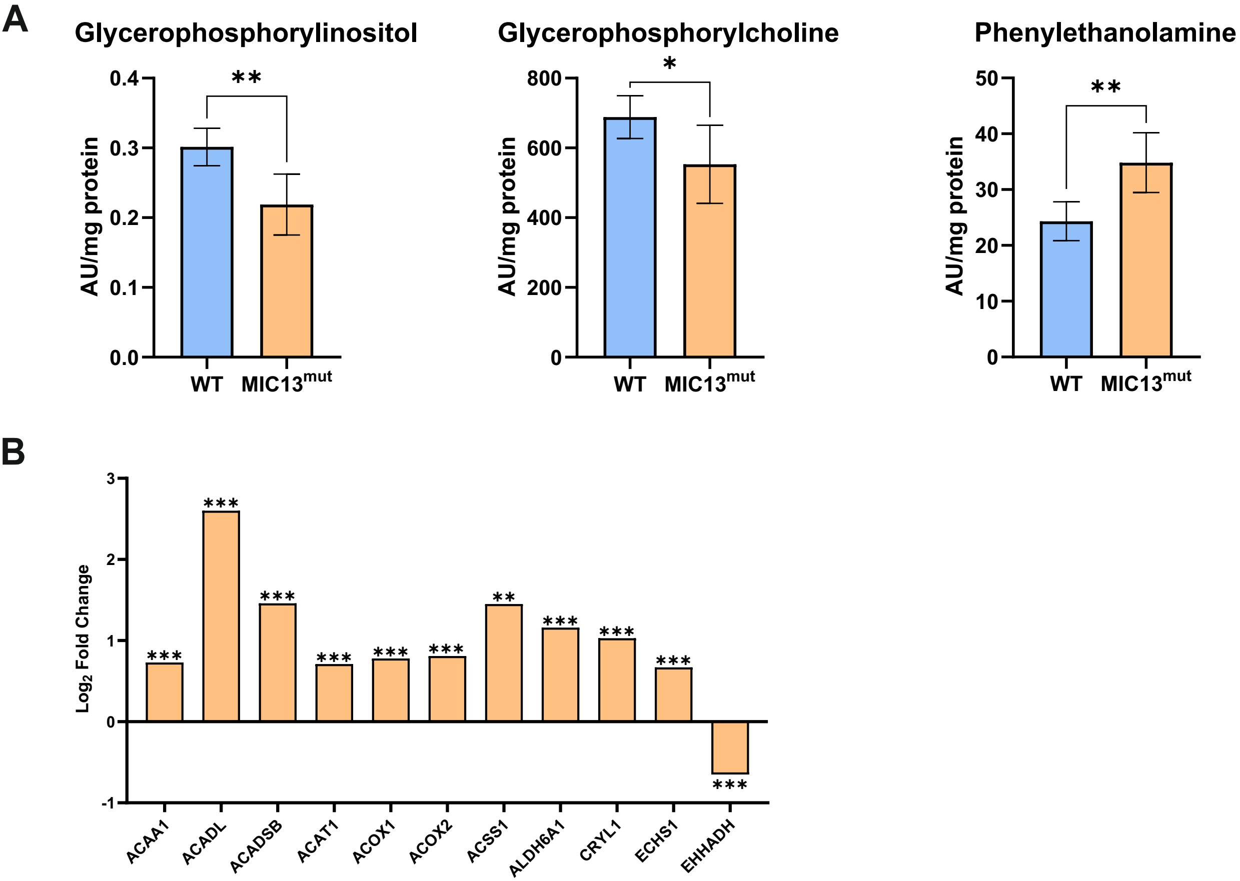

### Supplementary Figure 4

A

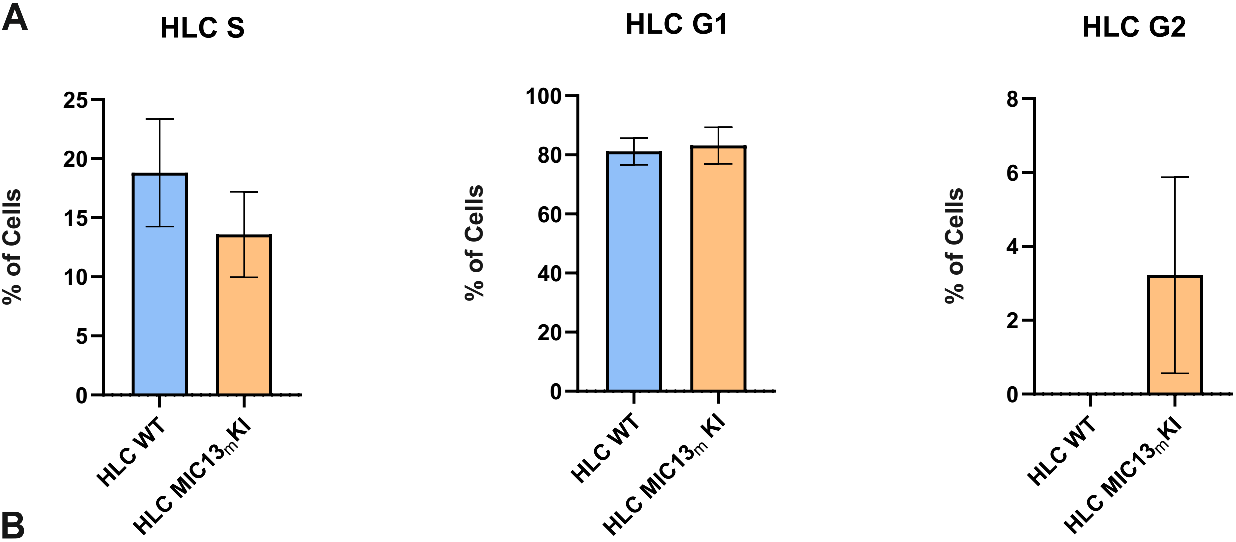


B

Gating strategy

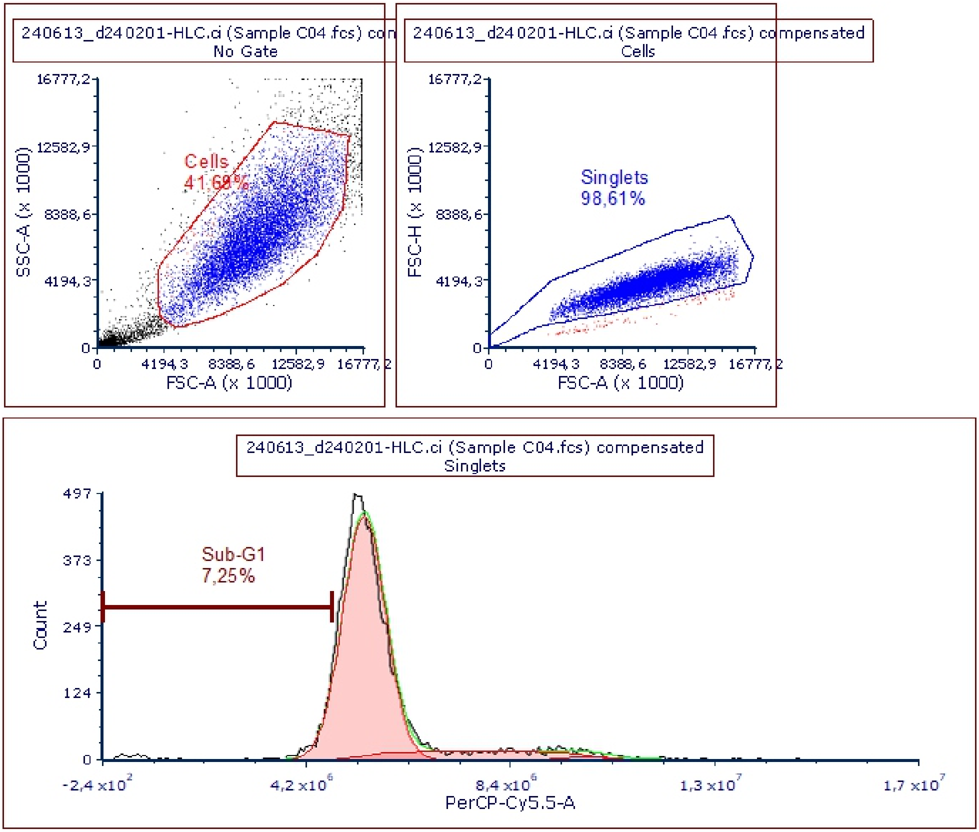
